# Diverging drivers of alpha and beta diversity across management types in Swedish boreal forests

**DOI:** 10.1101/2025.08.28.672851

**Authors:** Faith AM Jones, Alwin A Hardenbol, Albin A Larsson Ekstrom, Anne-Maarit A Hekkala, Mari Jonsson, Matti Koivula, Joachim Strenbom, Jorgen Sjogren

## Abstract

1. Managing multifunctional forest landscapes requires a better understanding of how biodiversity dimensions respond to habitat structures and management regimes. While species richness is commonly used to assess conservation value, it may not capture compositional differences critical for maintaining regional biodiversity.
2. We examined how local species richness (alpha diversity) and community turnover (beta diversity) relates to forest structure and management in Swedish boreal forests. We surveyed lichens, bryophytes, and polypore fungi, representing three sessile taxo-ecological groups across 120 sites.. The sites were distributed between three management types: young production forests, retention patches (trees left during harvest to support mature forest species in production forests), and unmanaged set-asides. We used generalised linear models and generalised dissimilarity modelling to assess how habitat structures (deadwood and living tree metrics) and spatial distance explained patterns of richness and turnover across taxa and management types.
3. For all taxa, species richness increase with lying deadwood volume. Lichen species richness also increased with living tree volume, and decreased with tree density in retention patches. Lichens had higher species richness in retention patches and set-asides, whereas only set-asides had statistically higher bryophyte richness, and there were no differences for polypore richness. In contrast, beta diversity was more often associated with geographic distance, with only limited influence of deadwood volume. The relationships between lichen beta diversity and habitat structures were not consistent between retention patches and set-asides, suggesting that retention patches may have limitations in conserving mature forest species despite supporting similar lichen species richness.
4. *Synthesis and applications*. Our results demonstrate that alpha and beta diversity can be driven by different ecological processes and management contexts, and that while tree retention can support higher biodiversity than young production forests in some taxa, it cannot replace the value of conserving mature forests. Effective conservation therefore needs to combine habitat quality and spatial complementarity when planning conservation initiatives to maintain diversity patterns across managed landscapes.

## 1. Intro

Balancing biodiversity conservation with other land use needs, such as carbon storage and production, is challenging and requires extensive ecosystem knowledge (Moilanen et al., 2011). In Europe, forests are a dominating land-cover type, but their conservation status remains mostly unfavourable (European Environment Agency, 2019). To support biodiversity and multifunctionality of forests, the EU Forest Strategy for 2030 proposes a Closer-to-Nature Forest Management strategy (European Commission: Directorate-General for Environment, 2023). This strategy emphasises the need to preserve biodiversity-rich habitats, as well as retaining or restoring important structures within areas managed for economic gain.

Boreal forests cover approximately 29% of the global forest area, and support a wide array of species and taxonomic groups (Kayes and Mallik, 2020). In boreal forests, many species are dependent on mature forest conditions, including stable microclimates and diverse substrates (Baker et al., 2016; Fossestøl and Sverdrup-Thygeson, 2009; Koivula and Vanha-Majamaa, 2020; Parajuli and Markwith, 2023; Yang et al., 2021; Zhang et al., 2024). The main boreal forest management system in Fennoscandia is a combination of retention forestry, where individual or groups of trees are retained during clearcutting, and set-asides, where forest areas are ad conserved for biodiversity purposes. This management system aims to emulate natural disturbance regimes in mature forests (Berglund and Kuuluvainen, 2021), and provides an opportunity for exploring how different management types can affect the relationship between biodiversity and habitat structures. There is significant variation, however, in how different taxa relate to forest structure (Kärvemo et al., 2021) and the effectiveness of retention forestry to conserve species (Baker et al., 2015).

Effective mixed-use conservation planning requires detailed knowledge of how biodiversity relates to habitat quality and availability. To this end, we assess biodiversity patterns in three representative management types in Swedish boreal landscapes: young forest stands (clearcut 20–30 years ago and managed for timber production), retention patches (small patches of forest left unharvested during the clearcutting of our young stands), and set-asides (larger areas of mature forest set-aside from harvesting activities to support biodiversity). Conditions in set-aside forests are closer to mature, closed-canopy forests; they have not recently experienced timber extraction activities and have been identified as key habitats of conservation value (Häkkilä et al., 2021; Timonen et al., 2011). Retention patches, though embedded within young forests, should maintain remnants of mature forest conditions, such as older trees and localised microclimate buffering, thereby supporting higher species richness than young forests, and offering temporary refugia for sensitive taxa (Baker et al., 2015; Perhans et al., 2009). In contrast, young production forests are often structurally homogeneous and lack key habitat features like large deadwood or mature canopy trees, making them less suitable for species dependent on complex, stable habitats (Hautala et al., 2011; Koivula and Vanha-Majamaa, 2020; c.f. Rudolphi and Gustafsson, 2011).

The links between biodiversity and forest structures have traditionally been quantified using local species richness (e.g Basile et al., 2021; Storch et al., 2023; Zeller et al., 2023), but many of these studies lack information on how species composition varies with habitat structure. Beta diversity, which quantifies how much species composition varies, describes a site’s complementary values for regional biodiversity. Beta diversity assessments allow us to consider trade-offs between protecting the areas with the highest local species richness and protecting the greatest variety of sites to maintain higher regional diversity. Incorporating beta diversity is therefore essential for protecting regional diversity (Socolar et al., 2016).

We focus on three sessile taxa that inhabit woody structures in the Swedish boreal forest: lichens growing on living and dead trees, bryophytes growing on lying deadwood, and polypores growing on standing and lying deadwood. Although all three taxa are sensitive to changes in forest conditions, their responses to human activities in boreal forests vary (Nirhamo et al., 2025). Lichens were less sensitive to felling activities than bryophytes (Hautala et al., 2011) and their responses to retention patch microclimate conditions was more variable (Perhans et al., 2009). Deadwood-inhabiting fungal communities shifted to more dehydration-tolerant fruiting structures in drier microclimates (Krah et al., 2022), but were also strongly influenced by dispersal processes (Norberg et al., 2019). There is therefore a need in conservation planning and management to explore variation in the relationships between species groups and the quantity of high-quality habitat structures. Doing otherwise risks missing important trade-offs in supporting different species when undertaking conservation initiatives.

First, we assess how alpha diversity (species richness) varies with forest structure across taxa and management types. Second, to explore trade-offs between species richness and compositional uniqueness, we use Generalised Dissimilarity Modelling (GDM)(Ferrier et al., 2007) to assess the relationship between beta diversity and site structures in each management type. We also include geographic distance between site-pairs in our GDMs. Sites are generally more similar in composition when they are closer geographically (Graco-Roza et al., 2022; Nekola and McGill, 2014; Soininen et al., 2007), but the strength and shape of this relationship holds ecological knowledge on connectivity, habitat similarity, and stochastic processes. Our main goal was to identify whether the same habitat structures drive both local species richness and compositional differences across management types. This integrated approach allows us to assess how forest structures interact with management contexts to influence different dimensions of biodiversity. In doing so, we provide applied insights into how conservation planning can move beyond richness hotspots to also capture compositional uniqueness and spatial complementarity in managed forest landscapes.

We have two main research questions, corresponding to alpha and beta diversity assessments:

1. How do key habitat structures (living trees and deadwood) influence species richness, and does that relationship vary with taxonomic group and management type?
2. What is the relative importance of differences in habitat structures compared to geographic distance in explaining species compositional variation, and how does that relationship vary with taxonomic group and management type?

Our three focal taxa differ in their ecological dependencies: lichens often colonise both living and dead trees, bryophytes often rely on moist deadwood substrates, and polypores largely depend on decomposing woody structures. We therefore expected their diversity patterns to respond differently to forest structure and management context. Specifically, we hypothesised that bryophyte and polypore diversity, as taxa strongly tied to deadwood (Dittrich et al., 2014; Hart et al., 2024; Sandström et al., 2019), would show stronger relationships with deadwood volume. Lichens diversity, conversely, would be more closely related to living tree volume, as their species richness usually increases with larger tree dimensions (Klein et al., 2020).

The relationship between species composition, habitat structures, and distance is less well studied. Distance is likely more important when it reflects larger-scale more heterogeneous underlying environmental or climatic gradients, land-use history, or biogeographic barriers across regions. Our forests are situated within the same forest habitat (EU biogeographic habitat: boreal forests) and country (Sweden), so we expected the differences in habitat features to influence beta diversity more then distance because differences in these features likely affect which niches are present at different sites.

Management type influences both habitat quality and continuity, shaping how taxa respond to structural features. Set-asides typically maintain mature forest characteristics like stable microclimates and high deadwood volumes, which benefit specialist and dispersal-limited species (Berglund and Kuuluvainen, 2021; Nirhamo et al., 2025). In contrast, young forests and retention patches may offer lower habitat quality and continuity, which may lead to weaker or more variable biodiversity–structure relationships, particularly for taxa reliant on old-growth conditions or complex substrates (Hautala et al., 2011; Perhans et al., 2009; c.f. Rudolphi et al., 2014). We therefore expected the species richness and beta diversity modelling to find strongest relationship between biodiversity and habitat structures in set-asides, and the weakest relationships in young forests.

## 2. Material and Methods

### 2.1 Study area

Our study area comprised 120 sites in Sweden, split equally between two regions within the same climactic zone, one in the east-centre of Sweden and the other in the south-west. Site distances varied from approximately 200 m to 330 km from each other.

We accounted for the distribution of forest management activities in our study regions by evenly selecting plots from three management types: young forests (harvested 20–30 years ago and representing typical intensive timber production forests in northern Sweden), retention patches (small patches of trees left unharvested in the young forests during original clearcutting activities), and set-asides (larger patches of forest unmanaged in recent decades to support biodiversity conservation).

All our forests had similar tree species compositions. They were dominated by Norway spruce (*Picea abies* (L.) Karst) and Scots pine (*Pinus sylvestris* L.) and contained some silver birch (*Betula pendula Roth.) and downy birch (B. pubescens* Ehrh.). There were also limited occurrences of other broadleaved trees including European aspen (*Populus tremula* L.), rowan (*Sorbus aucuparia* L.), goat willow (*Salix caprea* L.), and grey alder (*Alnus incana* (L.) Moench). The forest patches are located on mesic to moist soil types, avoiding the wettest and driest forests.

**Figure 1.**
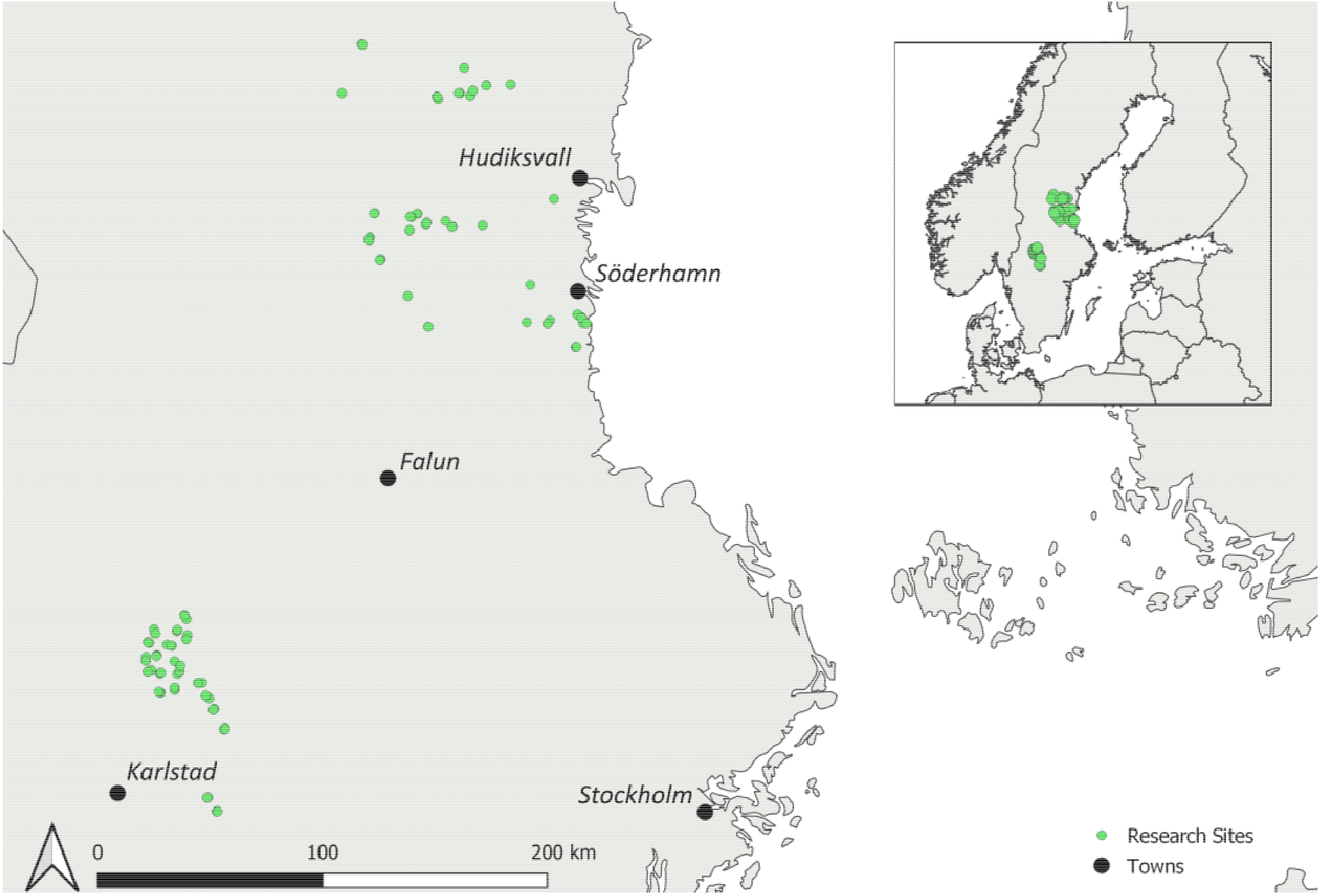
The locations of our 120 sites, distributed over two regions in Sweden. The three management types were distributed evenly across the research area. The inset contains a map of Fennoscandia showing the location of the sites relative to neighbouring countries. Many of the sites are too close together to show as separate points on the map so there are fewer than 120 points visible.

### 2.2 Species Data

We focus on three sessile, forest-dwelling taxa in this study. At site, we inventoried species from all three taxonomic groups in circular plots with a 20 m diameter. When retention patches were smaller than this, we surveyed smaller areas of down to 10 m diameter plots to avoid surveying non-retention patch habitat. We surveyed lichens on living and dead trees, bryophytes on lying deadwood, and polypores and a and a predefined list of corticioid fungi considered indicators of old-growth forests and of conservation concern (**Table S1**) on standing and lying deadwood. Full protocols are provided in the Supplementary Material.

### 2.3 Environmental variables

We surveyed a series of environmental variables at each of the 120 plots in conjunction with the species surveys. We focused on four structural features to reflect habitat quality and availability: the volume of standing deadwood, the volume of lying deadwood, the volume of living trees, and the density of living trees. The quantity of standing and lying deadwood may be indicators of high quality forest (Parajuli and Markwith, 2023) as they indicate conditions suitable for significant numbers of red-listed, deadwood-associated organisms. They also reflect habitat availability for bryophytes and polypores, and could influence whether deadwood-specialised lichens are present at a site. For living structure variables, we used the volume of living trees as a proxy for the amount of large trees on a site, and living tree stem density describes how open or closed the canopy is. Full details on survey protocos are provided in the Supplementary Material.

### 2.4 Species Richness Analysis

All statistics were undertaken in R version 4.2.2 (R Core Team, 2022). For assessing the relationship between lichen species richness and habitat structures, we used negative binomial generalised linear models using the *glm.nb* function of the R package *MASS*. We included a quadratic term for the relationship between lichen species richness and living tree volume, and checked it fitted better than a model without the quadratic term using AIC. For bryophytes and polypores, we used hurdle models using the *hurdle* function of the R package *pscl* (Zeileis et al., 2008) due to 19 sites without bryophytes and 34 without polypores (**Table S2**). These models fit two connected models: a presence-only model which provides information on which habitat structures relate to the probability of finding a species, and a species richness model for only the sites where species were present.

To capture as much variation in habitat availability and quality as possible, we included all four habitat structure variables in the models as well as management type. We also tested interaction effects between each management type and habitat structures but only included significant interactions in the final.

We standardised living tree density and living tree volume using the mean and standard deviation, whereas we log^10^ transformed lying and standing deadwood. We undertook post hoc comparisons using the *emmeans* function of the *emmeans* R package (Lenth, 2025) to test for significant differences between each management type where this variable was significant in the full model.

### 2.5 Generalised Dissimilarity Modelling (GDM) Analysis

We fit GDMs (Ferrier et al., 2007) to analyse the relationship between compositional turnover values of site pairs and the difference in geographic distance and environmental conditions. This method models the relationship between beta diversity of each site pair as a positive, non-linear function of geographic and environmental differences between these site pairs. The models calculate an I-spline function, called Partial Ecological Distance, describing the relationship between each explanatory variable and beta diversity keeping all other variables constant. The greater the sum of the Partial Ecological Distance curve, the higher the importance of the explanatory variable to explain beta diversity.

The response variable in each model was the Jaccard’s dissimilarity of each taxon in each management type. We used the *vegdist* function in the *vegan* R package (Oksanen et al., 2022) to calculate the beta diversity values for each site pair where a target taxon was found, giving a matrix of 7,140 beta diversity values for lichens (120 sites), 5,050 for bryophytes (101 sites), and 3,655 (84 sites) for polypores. We then used the *gdm* function to fit the GDM model and estimated the variable importance using the function *gdm.varImp*. To test the significance of each variable in each model we used the function *gdm.varImp*, which produces bootstrapped p values.

## 3. Results

In total, we identified 304 lichen, 149 bryophyte, and 66 polypore species across the 120 study sites. Environmental variables differed between management types. Set-asides were generally characterised by the highest standing and lying deadwood volumes, the highest volume of living trees and the lowest density of living stems (**Table S3**), meaning that they were stands with larger, older trees and highest volumes of both living and dead trees. Retention patches had similar values to set-asides, although they had lower variation between sites, and lower mean values of both living and dead tree volumes (**Table S3**). Young forests, conversely, had notably lower standing and lying deadwood volumes, lower living tree volumes, and higher living stem densities (**Table S3**), meaning they were young, dense living tree stands with few deadwood resources. None of the environmental variables correlated more than 50% with each other (**Table S4**).

### Species Richness

Lichens were found in all 120 sites, and their species richness varied with management type and environmental variables. Their species richness was higher in retention patches and set-asides than in young forests (**Table 1, Figure 2a, Table S5**) and increased with lying deadwood volume (**Table 1, Figure 2b**). Lichen species richness had a non-linear relationship with the volume of living trees, where species richness peaked with intermediate volume (**Table 1, Figure 2c**). Lichen species richness also decreased with increasing density of living trees in retention patches (**Table 1, Figure 2d**).

**Table 1.** The relationship between environmental variables and species richness from either negative binomial (lichens) or hurdle models with negative binomial (bryophytes and polypores). Standard errors are shown in the SE column, and P-values in the Pr column. Coefficients from the species presence part of the hurdle models are shown in **Table S7**. For lichens and bryophytes, where management type was a significant predictor of species richness in the models, post hoc comparisons between each management type are in **Table S5**. We only retained significant interactions for the final models. For the bryophyte model, the significant interactions were in the species presence part of the model (**Table S7**).

|  | Environmental Variables | Species Richness<br>Relationship | SE | Pr |
| --- | --- | --- | --- | --- |
| *Lichens* | Standing Deadwood Volume | 0.025 | 0.02 | 0.27 |
|  | Lying Deadwood Volume | **0.04** | **0.02** | **0.03** |
|  | Living Tree Volume | **-0.03** | **0.02** | **0.03** |
|  | Living Tree Density | 0.06 | 0.03 | 0.051 |
|  | Retention Patch | **0.4** | **0.06** | **<0.001** |
|  | Set-aside | **0.56** | **0.09** | **<0.001** |
|  | Living Tree Density : Retention<br>Patch | **-0.14** | **0.05** | **0.005** |
|  | Living Tree Density :<br>Set-aside | 0.06 | 0.08 | 0.44 |
| *Bryophytes* | Standing Deadwood Volume | -0.02 | 0.13 | 0.86 |
|  | **Lying Deadwood Volume** | **0.81** | **0.11** | **<0.001** |
|  | Living Tree Volume | 0.002 | 0.05 | 0.97 |
|  | Living Tree Density | 0.08 | 0.05 | 0.09 |
|  | Retention Patch | 0.11 | 0.15 | 0.44 |
|  | **Set-aside** | **0.57** | **0.16** | **<0.001** |
| *Polypores* | Standing Deadwood Volume | 0.16 | 0.17 | 0.34 |
|  | **Lying Deadwood Volume** | **1.10** | **0.15** | **<0.001** |
|  | Living Tree Volume | 0.005 | 0.06 | 0.93 |
|  | Living Tree Density | -0.08 | 0.10 | 0.40 |
|  | Retention Patch | -0.10 | 0.25 | 0.68 |
|  | Set-aside | 0.09 | 0.27 | 0.72 |

**Figure 2.**
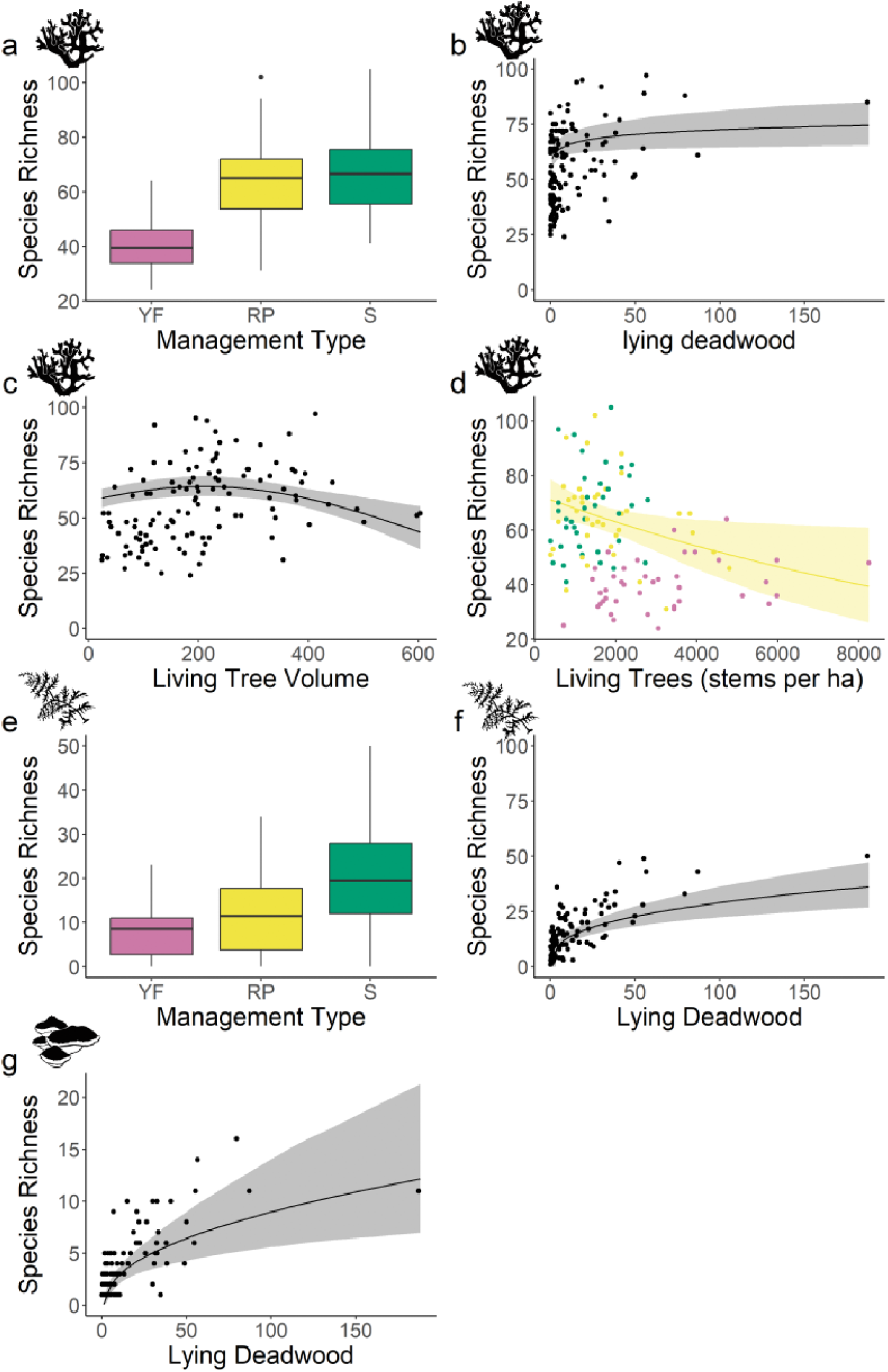
The significant relationships between environmental variables and species richness of lichens, bryophytes, and polypores. All other variables in the model were at their median values for plotting and, unless otherwise stated, the management type was Retention Patch. In panel (a) we show the distribution of lichen species richness values in each management type (YF = Young forest, RP = Retention patch, S = Set-aside). We show the relationship between lichen species richness and lying deadwood (b) and living tree volume (c) regardless of management type. Panel (d) shows the relationship between lichen species richness and living tree density. Points are coloured using the same key as the plot a, but there is only a yellow ribbon representing the significant relationship between lichen species richness in retention patches and living tree density (Table 2). There are no corresponding ribbons showing Set-asides or Young forests as there were no significant relationships between lichen species richness and living tree density in the models. In panel (e) we show the distribution of bryophyte species richness values in each management type (YF = Young forest, RP = Retention patch, S = Set-aside), and the relationship between bryophyte species richness and lying deadwood regardless of management type is shown in panel (f). Panel (g) shows the relationship between polypore species richness and lying deadwood, regardless of management type. For continuous relationships, the bold line shows the average relationship predicted by the model, and the ribbon represents the 95% confidence intervals around this model prediction. For boxplots, we show the median as the horizontal line in the box, the 75% and 25% quantile values as the upper and lower boundaries of the box, and the minimum and maximum values as the end of the whiskers.

Bryophytes were found in 101 of the 120 sites (**Table S1**), and varied between management types and with environmental variables (Table S6). The probability of finding at least one bryophyte species increased with the volume of lying deadwood (**Table S7**). For the sites with at least one bryophyte present, species richness was higher in set-asides than in retention patches or young forests **(Table 1, Figure 2e Table S5**) and increased with the volume of lying deadwood (**Table 1, Figure 2f**).

**Table 2.**
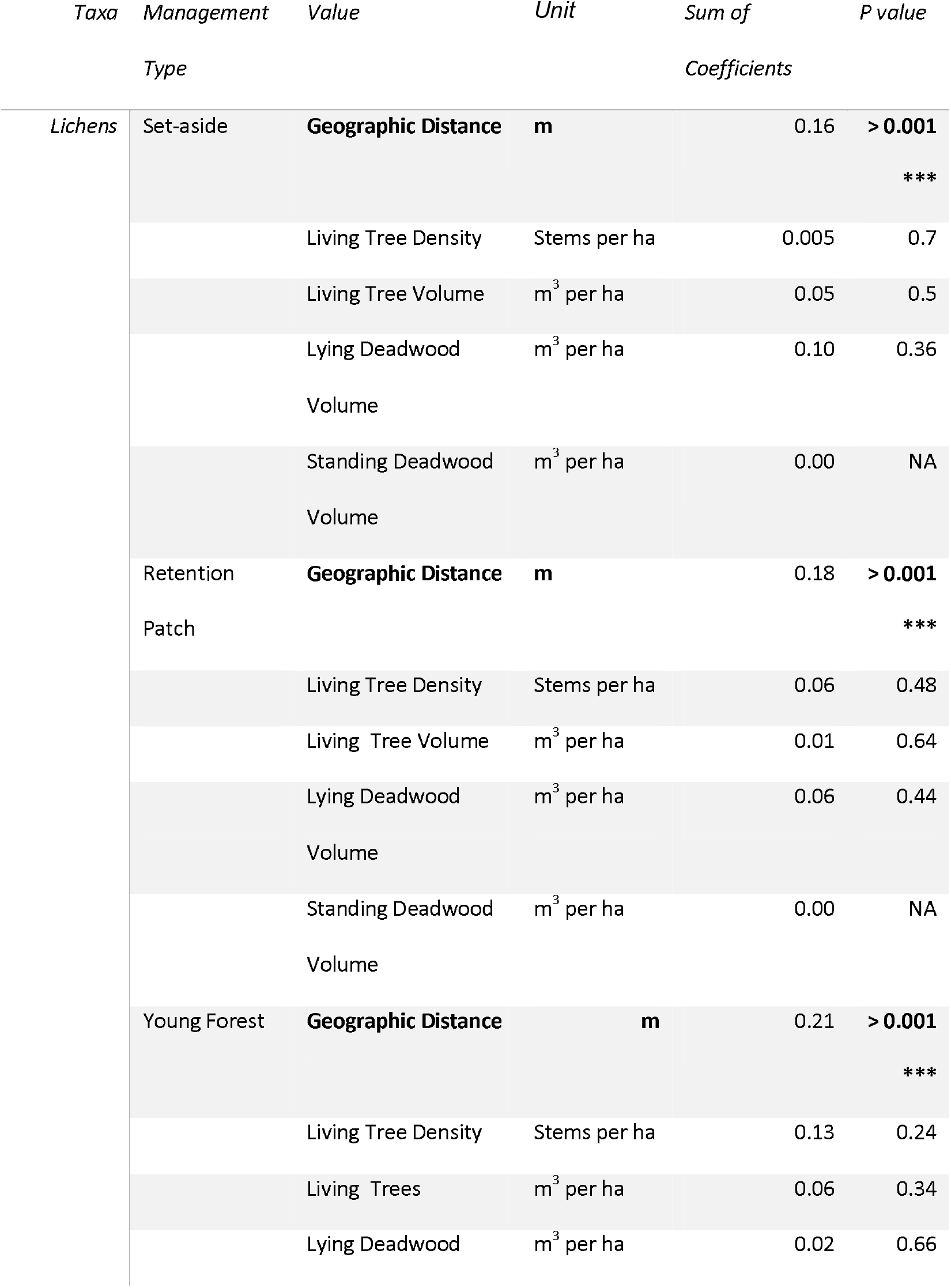
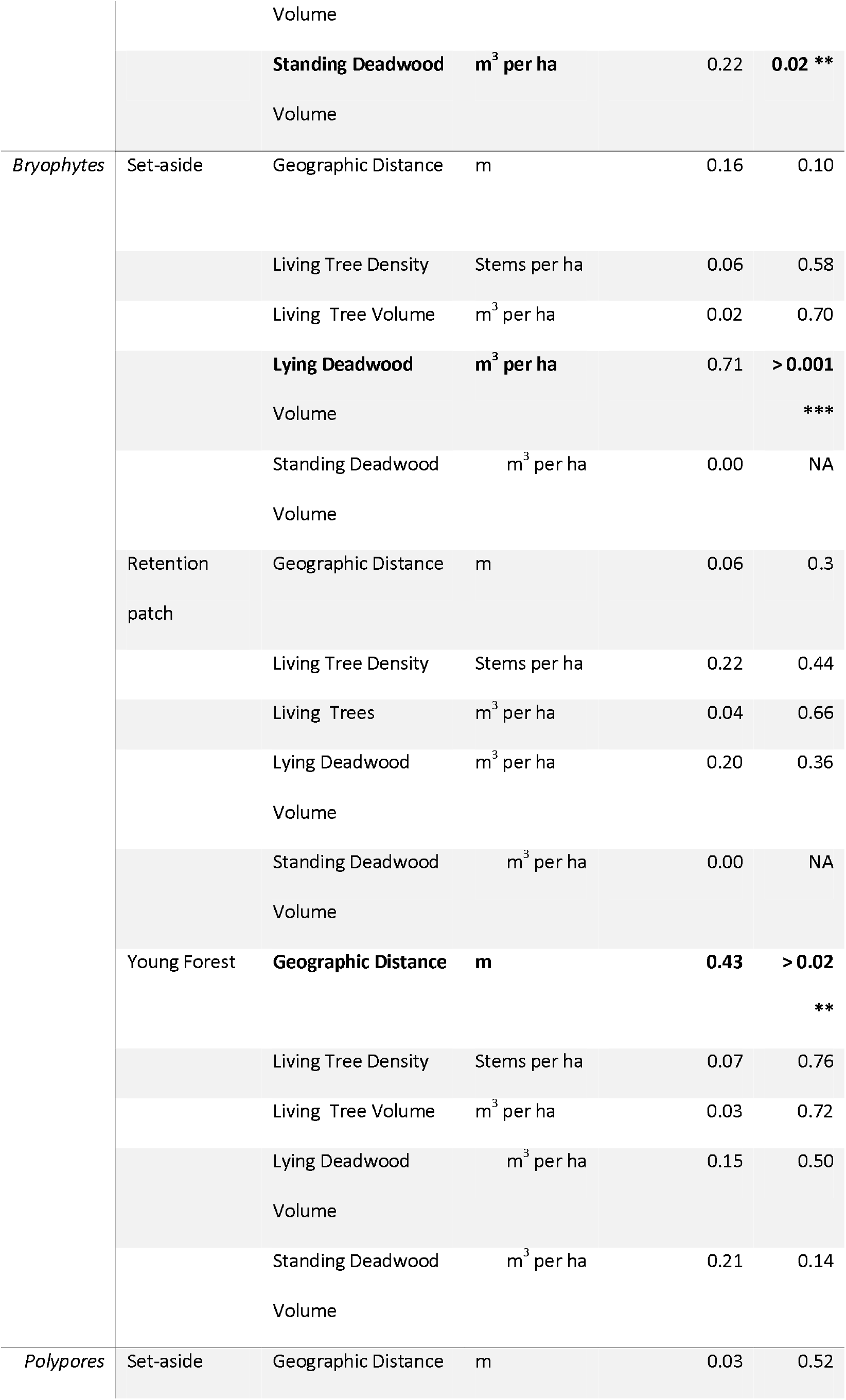
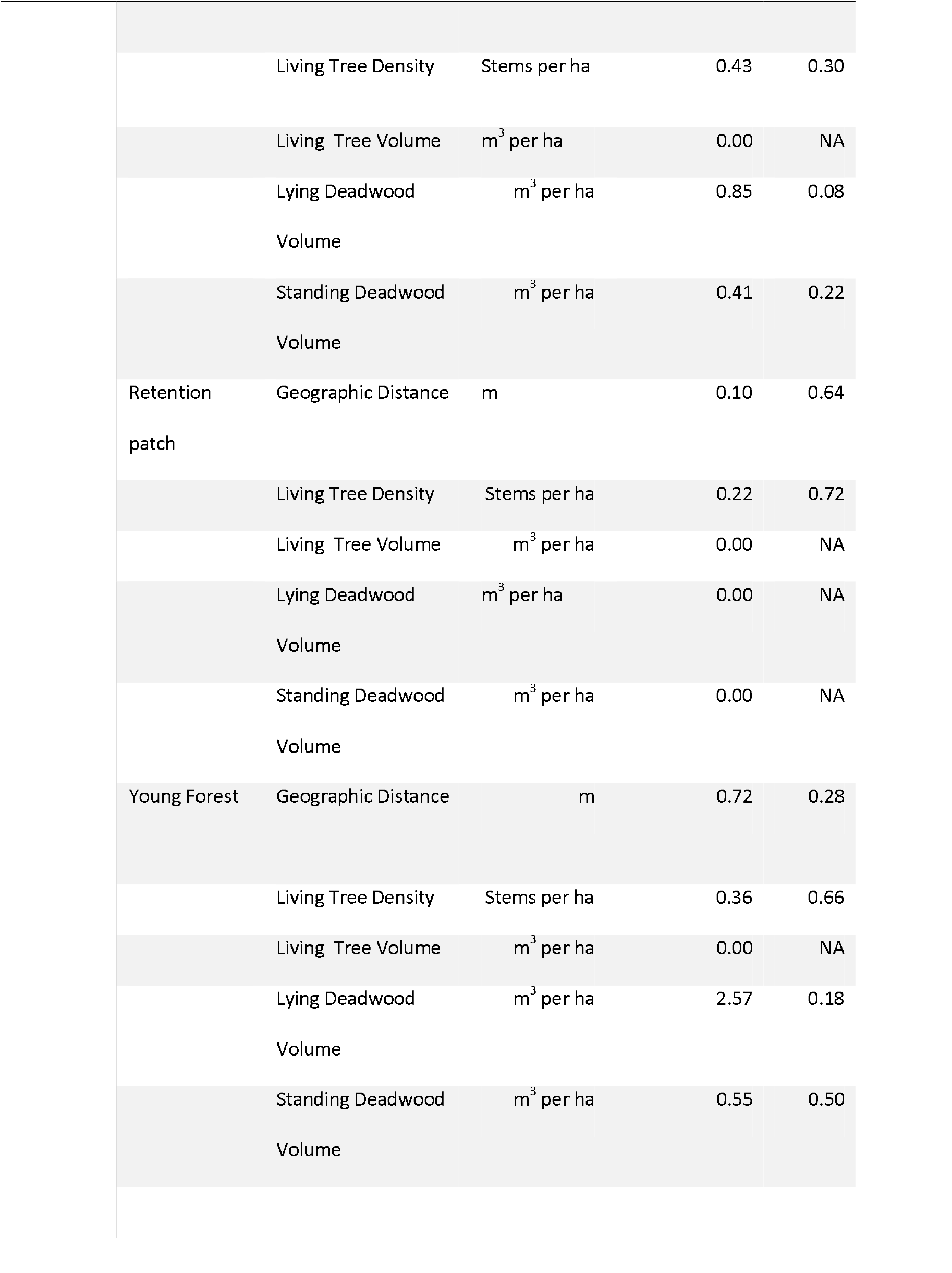
Generalised dissimilarity modelling (GDM) sum of I-spline coefficient values in explaining beta diversity patterns for each taxon and explanatory value combination, and associated significance of each value in the model. For the sum of I-spline coefficients, a larger value indicates a greater contribution of that variable to explaining beta diversity, and a smaller or near-zero value indicates little or no explanatory power for compositional variation. Significance was calculated using matrix permutation.

We found polypores in 84 of the sites (see **Table S8** for environmental variables), with species found more often in set-asides and least often in young forests (**Table S2**). The probability of finding at least one polypore species increased with the volume of lying deadwood (**Table S7**). At sites where at least one polypore species was recorded, species richness increased with increasing volume of lying deadwood (**Table 1, Figure 2g**). We found no effect of management type on polypore species richness.

### Beta diversity

Geographic distance was the only significant predictor of lichen beta diversity in both set-asides and retention patches, with compositional dissimilarity increasing with between-patch distance (**Table 2, Figure 3a&b**). The lichen Set-aside GDM included geographic distance and all four environmental variables as predictors, and explained 9.5% of the variation in observed pairwise dissimilarities with an intercept of 0.40. The lichen Retention Patch GDM had the same structure, and explained 11.1% of the variation in observed dissimilarities, with an intercept of 0.34. Although geographic distance, volume of lying deadwood, volume of living trees, and density of living trees all had sums of coefficients greater than zero in both models (**Table 2, Figure S2 & Figure S3**), and geographic distance was the only significant variable (**Table 2, Figure 3a&b**). In young forests, lichen beta diversity increased significantly with both geographic distance and difference in standing deadwood volume (**Table 2, Figure 3 & d**). The lichen Young Forest GDM explained 30% of the variation in pairwise dissimilarities and had an intercept of 0.30. Geographic distance and all four environmental variables had sums of coefficients greater than zero (**Figures S4**), but only geographic distance (**Table 2, Figure 3c**) and standing deadwood had significant positive relationships with increasing lichen beta diversity.

**Figure 3.**
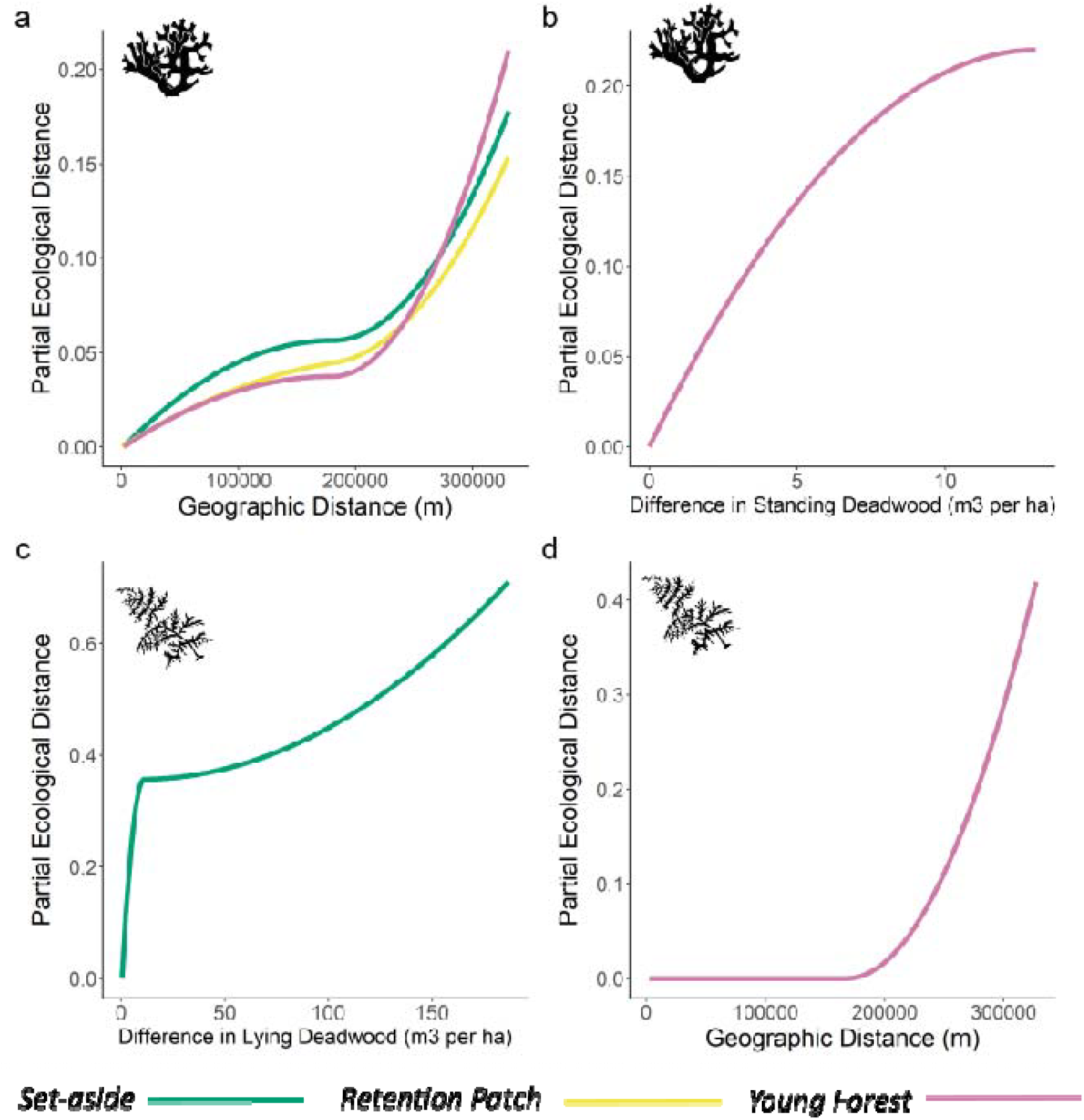
Generalised dissimilarity modelling (GDM) I-spline partial functions (partial ecological distance), showing the rate of species turnover along the significant geographic and environmental gradients of each taxon–management type model combination. Partial Ecological Distance represents the beta diversity along the specific habitat or geographic gradient, holding all other variables constant. A steeper curve means that beta diversity changes more rapidly with differences in the x-variable. For lichen models, geographic distance was significant in the Set-aside, Retention Patch, and Young Forest models (a). In each case, the curve was shallower when sites were close together, but much steeper when sites were at least 181 311 m, 179 379m or 179 453 m apart, respectively. Standing deadwood was only significant in the Young Forest (b) model and showed a stronger positive relationship with beta diversity when differences in standing deadwood were smaller. For bryophytes, the only significant variables were lying deadwood in the Set-aside model (c), where partial ecological distance changed abruptly with differences of between 0 and 11m^3^ per ha in lying deadwood, and Geographic Distance in the young forest model (d), where sites were very similar within a 181 311 m distance but very different after this point. There were no significant variables in the bryophyte Retention Patch model, nor any of the polypore models. Complete plots of all I-spline functions with a sum of coefficients greater than zero can be found in **Figures S2–S8**..

In set-asides, bryophyte beta diversity increased with differences in lying deadwood, especially with when differences were below 11 m^3^ per ha (**Table 2, Figure 3e**). The bryophyte Set-aside GDM explained 28.8% of the variation in observed dissimilarities, with an intercept of 0.56. Although geographic distance, density of living trees, volume of living trees, and volume of lying deadwood all had sums of coefficients greater than zero (**Table 2, Figure S5**), only lying deadwood significantly influenced the model output (**Table 2, Figure 3e**). We found no relationship between either geographic or environmental variables and bryophyte beta diversity in retention patches (**Table 2**). The bryophyte Retention Patch GDM explained only 3.6% of the variation in the model, and had an intercept of 0.69. Geographic distance, density of living trees, volume of living trees, and volume of lying deadwood all had sums of coefficients greater than zero (**Figure S6**), but none were significant (**Table 2**). In young forests, beta diversity increased significantly with geographic distance, but only once distance between sites was above approximately 200,000 m. The bryophyte Young Forest GDM explained 9.2% of the variation in the model, and had an intercept of 0.66. Geographic distance and all four environmental variables had sums of coefficients greater than zero (**Figure S7**), but only geographic distance was significant (**Table 2, Figure 3f**),

We found no relationships between polypore beta diversity and either geographic or environmental differences in any management type (**Table 2**). The polypore Set-aside GDM explained 12.5%, with an intercept of 0.98. Geographic distance, density of living trees, volume of lying deadwood, and volume of standing deadwood all had coefficients summing to more than zero (**Figure S8**), but none were significant (**Table 2**). The polypore Retention Patch GDM explained only 0.23% of the variation in the observed dissimilarities in the model, and had an intercept of 1.8, signalling a fit with very low information from the explanatory variables. Only geographic distance and density of living trees had coefficients greater than zero (**Figure S9**), and none were significant (**Table 2**). The polypore Young Forest GDM explained 10% of the variation in observed dissimilarities and had an intercept of 1.7. As with the polypore Set-aside model, geographic distance, density of living trees, volume of lying deadwood, and volume of standing deadwood all had coefficients summing to more than zero (**Figure S10**), but none were significant (**Table 2**).

## Discussion

We tested the assumption that different aspects of biodiversity relate similarly to the same habitat structures, implicit in analysis using only species richness for conservation planning. We showed that species richness and species composition (beta diversity) often responded differently to environmental variables. Species richness was most often related to either the density of living trees or the volume of lying deadwood, although the strength of these relationships depended on species groups and management type. Lichen and bryophyte beta diversity related to geographic distance. Unlike the species richness, lichen beta diversity related to standing deadwood rather than lying deadwood. Bryophyte beta diversity related to the volume of lying deadwood alongside geographic distance. We found no variables relating to the beta diversity of polypores.

### Contrasts between alpha and beta diversity

We found evidence that while species richness responded to habitat quantity and quality (e.g., the volumes of living and dead trees, and density of living trees), beta diversity patterns were less predictable and more commonly influenced by the spatial distance and management type. Lying deadwood and living tree volumes explained lichen species richness, whereas, beta diversity was more closely aligned to spatial distributions and, to a lesser extent, the volume of standing deadwood. For bryophytes, larger differences in the volume of deadwood between management types also correlated with larger differences in the compositions of compared species communities. Polypore species richness was significantly higher in sites with more lying deadwood, but no variables explained patterns of polypore beta diversity. The contrasting patterns we found between alpha and beta diversity assessments highlight the limitations of conservation strategies focused solely on maximising species richness (Luby et al., 2022). Our results demonstrate that, although it is often poorly represented in applied monitoring and planning (Mokany et al., 2022; Socolar et al., 2016), conservation planning needs to explicitly account for beta diversity to protect maximum biodiversity variation rather than extrapolating from species richness patterns.

Our results on beta diversity emphasise the importance of considering the spatial context in lichen conservation planning (Gustafsson et al., 2025; Williams and Ellis, 2018). The influence of geographic distance increased substantially once sites were around 180 km apart, which likely reflects a shift from within-region to between-region comparisons. Although both regions are in the same climactic zone and habitat type (boreal forest), this differentiation may be because they have experienced different intensities of historical forest activities (Lundmark et al., 2013). Additionally, limited dispersal ability, particularly for bryophytes and lichens with poor potential for long-distance dispersal (Sillett et al., 2000), could lead to spatial turnover.

### Taxon-specific responses

Retention patches harboured similar lichen species richness to set-asides, and more species than young forests. This result supports the effectiveness of retention forestry at providing refugia for many lichen species (Perhans et al., 2009; Rudolphi et al., 2014). We also found a negative relationship between tree-stem density and lichen species richness in retention patches. Our results are in line with previous evidence that increased shading from dense forests decreases the species richness of epiphytic lichens (Klein et al., 2020). One possible explanation for the lack of relationship between the species richness of lichens and the density of living trees in the other management types is low variation in stem density in these management types: young forests had high stem densities typical of dense young production stands, whereas the stem densities in set-asides were relatively low, as in larger, more mature forests. In addition, we only found relationships between the volume of standing deadwood and beta diversity in young forests.

Geographic distance showed the same relationship to lichen beta diversity in all management types, suggesting that spatial processes are affecting the different forest communities in similar ways. In contrast, differences in standing deadwood only related to lichen beta diversity in young forests. Beta diversity increased strongly with increasing standing deadwood up to approximately 10 m^3^ per ha, and then the relationship levelled off. Standing deadwood is an important habitat lignicolous and corticolous lichen abundance and diversity in managed forests (Santaniello et al., 2017; Svensson et al., 2016). Epiphytic lichen species do not easily disperse long distances (Dettki et al., 2000), so groups of similar habitat structures closer together are important for maintaining their diversity (Klein et al., 2021). In young production forests with little standing deadwood, lichen communities may be dominated by generalists that use both living and dead trees, and by species with greater dispersal ability. A previous study has shown that retaining old-growth structures, such as old living and dead trees, in young forests after final harvest is important for lichens (Rudolphi et al., 2014), and our results corroborate this finding. Thus, to support lichen biodiversity also in boreal young forests, more standing deadwood should be maintained and encouraged during production management activities.

Bryophyte species richness was similar between retention patches and young forests, suggesting retention patches are less effective for bryophytes than lichens. An explanation for this result is that the retention patches may be too small to function as effective refugia for the more sensitive bryophyte species, where lichen species richness may be bolstered by more robust species (Perhans et al., 2009). Bryophytes may require buffers of up to 30m from south-facing edges to conserve microclimatically sensitive species, an edge size rarely available in typical Swedish retention patches (Löbel et al., 2012). However, this conclusion may contrast with previous evidence that sensitive bryophytes can survive to a large extent in retention patches (Rudolphi et al., 2014). In our study, we did not categorise habitat preferences or sensitivities of the species detected in the different forest types, though, so we cannot know if this contrast is due to an overabundance of less sensitive bryophytes dominating our results.

While bryophyte species richness increased with lying deadwood volume, bryophyte species composition was only related to this environmental variable in set-aside forests. This result suggests that an increase in deadwood will lead to distinct and larger species communities only in the more natural set-aside forests. An explanation for the disconnect between bryophyte species richness and composition is that more heavily managed forests do not have sufficient amounts of deadwood to support unique communities, or that deadwood quality is higher in mature, set-aside forests. The type and variety of deadwood may be more important to deadwood-inhabiting species than deadwood volume (Parajuli and Markwith, 2023)(Sandström et al., 2019), for example different decay stages, size classes, and tree species. Deadwood volume may be a proxy for deadwood quality in set-asides with more natural deadwood accumulation and more tree species to provide deadwood, but may be decoupled from deadwood volume in other management types where deadwood may be predominantly single-aged spruce and pine. Another possibility is that conditions in young forests and retention patches are less favourable for the types of species found in high-deadwood set-asides due to factors we did not include in our model, for example historical management intensity (Táborská et al., 2017) or microclimate (Baker et al., 2016). Regardless, our results for bryophytes emphases the importance of conserving high quality mature forests within managed forest landscapes to help maintain regional species pools.

We found higher polypore richness with more lying deadwood, in line with previous studies, (Sandström et al., 2019; Ylisirniö et al., 2016), highlighting habitat availability as a key driverWhile this may reflect the species–area relationship (Scheiner, 2003), richness did not increase with standing deadwood, suggesting other factors such as deadwood age and quality (Parajuli and Markwith, 2023; Sandström et al., 2019). Because standing deadwood is often more recent, polypores may not have fully colonised it compared to lying deadwood.

Surprisingly, we found no relationship between polypore beta diversity and any of our habitat or geographic distance variables. Deadwood-inhabiting polypore communities are known to exhibit very high beta diversity and turnover in managed forests (Halme et al., 2013), so this result may be due generally high turnover between sites overwhelming differences in habitat characteristics. Their community assembly is a highly stochastic process, even with detailed descriptions of substrate characteristics (Norberg et al., 2019). Polypore GDMs had a low explanatory power of 10%–15%, far below the 20%–50% found in most ecological models (Mokany et al., 2022). By including different site-level habitat variables, for example, microclimate variation (Ylisirniö et al., 2016), host tree diversity (Renvall, 1995), or deadwood decomposition (Brūmelis et al., 2017) and diversity (Yang et al., 2021), we could potentially improve the GDM’s power. We also only surveyed visible fruiting bodies, so may have missed species present in the deadwood that were not fruiting at the time. However, we undertook surveys at peak fruiting season to minimise this issue, and fruiting-body-based community surveys can give comparable results to DNA-based surveys (Saine et al., 2020), as well as yield species–environmental relationships.

### Conservation implications

Our results offer actionable insights for advancing the EU’s Closer-to-Nature Forest Management approach, which promotes sustainable forest production though greater integration of tree retention and setting aside forests for biodiversity conservation (European Commission: Directorate-General for Environment, 2023). Our results suggest that retention forestry does not fully replicate the biodiversity–habitat relationships found in mature forests. We found lower species richness, especially in young forests, suggesting limited colonization or persistence of species. In addition, relationships between beta diversity and forest structures were inconsistent between management types, suggesting a failure of retention forestry to mimic the natural processes linking species to their environment in mature forests. These results imply that retention forestry does not fully substitute for mature forest conditions (Berglund and Kuuluvainen, 2021; Klein et al., 2020). Retention patches, which should act as refugia for sensitive mature-forest species, are often too small to provide more than low quality edge habitats (Curzon et al., 2020; Perhans et al., 2009). There is mixed evidence for the effectiveness of retention trees to support red-listed lichen species. In some studies there were high survival rates of these sensitive lichen species(e.g. Gustafsson et al., 2013; Rudolphi et al., 2014). In contrast, in another study - individual trees retained in young forest failed to support target old-growth epihphyic species due to exposure and risk of windthrow (Löbel et al., 2012). We found that even small increases in standing deadwood volume in young forests make a positive contribution to maintaining lichen species pools, supporting the role of retaining trees during in sustainable forest management. However, overreliance on retention forestry is unlikely to support boreal forest biodiversity (Roberge et al., 2015) as retention forestry cannot replace the value of protecting larger areas of mature forest (Gustafsson et al., 2025).

Given that beta diversity was more strongly structured by geographic distance than habitat variables, particularly for lichens, our findings demonstrate the importance of spatially explicit conservation planning. This result supports calls to design forest conservation networks that ensure broad spatial representation across landscapes to capture regional variation in species composition (Mokany et al., 2022; Socolar et al., 2016). Focusing only on areas of high local richness risks overlooking rare or complementary communities, especially for dispersal-limited taxa (Sillett et al., 2000; Williams and Ellis, 2018). As we demonstrate, GDMs show promise as a useful tool to facilitate incorporating species composition into conservation planning (Mokany et al., 2022). This means that Closer-to-Nature Forest Management implementation should go beyond enhancing structural elements within production forests. Forest planning should integrate mature set-asides distributed across the landscape, complemented by retention patches designed to maintain habitat heterogeneity.

Sweden’s boreal forests have experienced significant economic exploitation for centuries (Kayes and Mallik, 2020). Even the most natural set-aside forests have experienced some degradation from human activities (Lundmark et al., 2013). The amounts of deadwood in our study are in line with the Swedish national average of 7.6 m^3^ per ha (Jonsson et al., 2016) but often fall far below the 20 m^3^ per ha suggested for, for example, effective polypore species conservation (Ylisirniö et al., 2016). Old growth forests host a unique selection of species not found in the more intensively managed forests included in our study (Nirhamo et al., 2025), and unaccounted for in our assessment. Restoration could also support increased structural diversity for a variety of taxa in more heavily managed forests (Espinosa Del Alba et al., 2021), and may be necessary to support these old-growth species in a forest landscape currently lacking in old-growth quality habitats (Hjältén et al., 2023).

## Conclusion

Our findings demonstrate a decoupling between species richness and species composition across taxa in boreal landscapes, underscoring the importance of evaluating multiple biodiversity dimensions for effective conservation planning (Socolar et al., 2016). While structural habitat features such as deadwood and living tree volume influenced species richness, beta diversity patterns were more strongly shaped by geographic distance and, particularly in more natural forests. On a global scale, species richness prioritisation (e.g. Luby et al., 2022) has merit for highlighting regions at risk of losing the most species. Our results, though, emphasise the need to incorporate beta diversity into spatial planning frameworks to maintain ecological heterogeneity and avoid homogenisation of forest habitats (Berglund and Kuuluvainen, 2021; Kärvemo et al., 2021; Koivula and Vanha-Majamaa, 2020)

