## Supplementary Material for "Diverging drivers of alpha and beta diversity across management types in Swedish boreal forests"

for

### Full Species survey protocols

We surveyed lichens on all standing dead and a subset of living trees, recording occurrences up to 2 m from the ground. The subsample consisted of 10 trees per plot: three small trees (diameter at breast height (DBH) of 5–15 cm) and seven large trees (DBH of at least 15 cm) of the dominant tree species. If an insufficient number of trees was available in one size class, we surveyed more from the other size class to maintain a constant subsample of 10 living trees per plot. Most lichens were identified to species in the field, but some difficult-to-identify specimens were further assessed in the laboratory. We used the function *DataInfobeta3D* from the R package *iNEXT.beta3D* (Chao et al., 2023) to ensure all samples reached at least 90% coverage of species present despite subsampling living trees.

For the bryophytes, we surveyed all lying deadwood with a minimum diameter greater than 15 cm and an additional five smaller deadwood items per dominant tree species. Highly decayed wood (Decay stage 5, Renvall, 1995) was not surveyed as we considered it part of the organic top soil layer.

For the polypore survey, we surveyed deadwood-inhabiting polypores and a predefined list of corticioid fungi considered indicators of old-growth forests and of conservation concern (**Table S1**). We surveyed three small lying items of deadwood (DBH of 5–15 cm) and three small standing trees (DBH of 5–15 cm) per dominant tree species. We also surveyed all deadwood with either a DBH of at least 15 cm (for standing deadwood) or a minimum diameter of 15 cm and a minimum length of 1.3 m (lying deadwood) for all tree species including broadleaves. Standing trees were surveyed up to a height of 2 m. Polypore species were identified based on the appearance of fruiting bodies.

Table S1. Corticioid species that serve as old-growth forest indicators or that are of special conservation interest in boreal forests that were surveyed as part of the polypore surveys.

| **Tree Species** | **Corticioid species** |
| --- | --- |
| *Picea abies* | *Asterodon ferruginosus*  *Conferticium ochraceum*  *Cystostereum murrayi*  *Laurilia sulcata* (prob more northern sp) - taigaskinn  *Phlebia centrifuga* |
| *Pinus sylvestris* | *Chaetodermella luna*  *Crustoderma corneum*  *Crustoderma dryinum*  *Irpicodon pendulus -* vintertagging  *Odonticium romellii*  *Pseudomerulius aureus* |
| Additional *Pinus sylvestris* spp included in Finland but NOT Sweden, mykorrhiza or not “signalart”: | *Hydnellum gracilipes*  *Phlebia serialis*  *Sistotremastrum suecicum*  *Sparassis crispa* |
| Broadleaves (mainly *Populus*, *Salix*, *Betula* NOT *Ulmus*, *Quercus*, *Fagus*, *Fraxinus* etc) | *Hydnocristella himantia* |

### Full Structure survey protocols

Deadwood was surveyed in the full 20-m radius plots, and was categorised into four types: whole lying trees, broken lying trees, standing whole trees, and standing broken trees. All deadwood items with a mean diameter or estimated Diameter at Breast Height (DBH, 1.3 m from the bottom of a standing or lying whole tree) of at least 10 cm within the species survey plots were recorded. For whole lying and standing dead trees, we measured DBH and used region specific allometric equations to estimate volume (Equation S1). For broken standing deadwood we also used the DBH and allometric equations, but detracted the volume based on the height at which the tree was broken. For lying broken trees, we estimated volume using length and either mean or minimum and maximum diameter measurements.

For living tree data, we surveyed all living trees taller than 1.3 m within a 7-m radius plot centred on the middle of the 20-m radius species survey plots. We recorded DBH and tree species, and then sed allometric equations, calibrated to each region, to estimate volume (Ollas, Rune., 1980 , Equation S1). All variables were converted to per ha values before analysis.

### Tree Volume Calculations

Equation S1. The formulae used to estimate tree height and volume of spruce, spine and broadleaf trees in our two regions, Halsingland (northern region) and Varmland (south west regions).

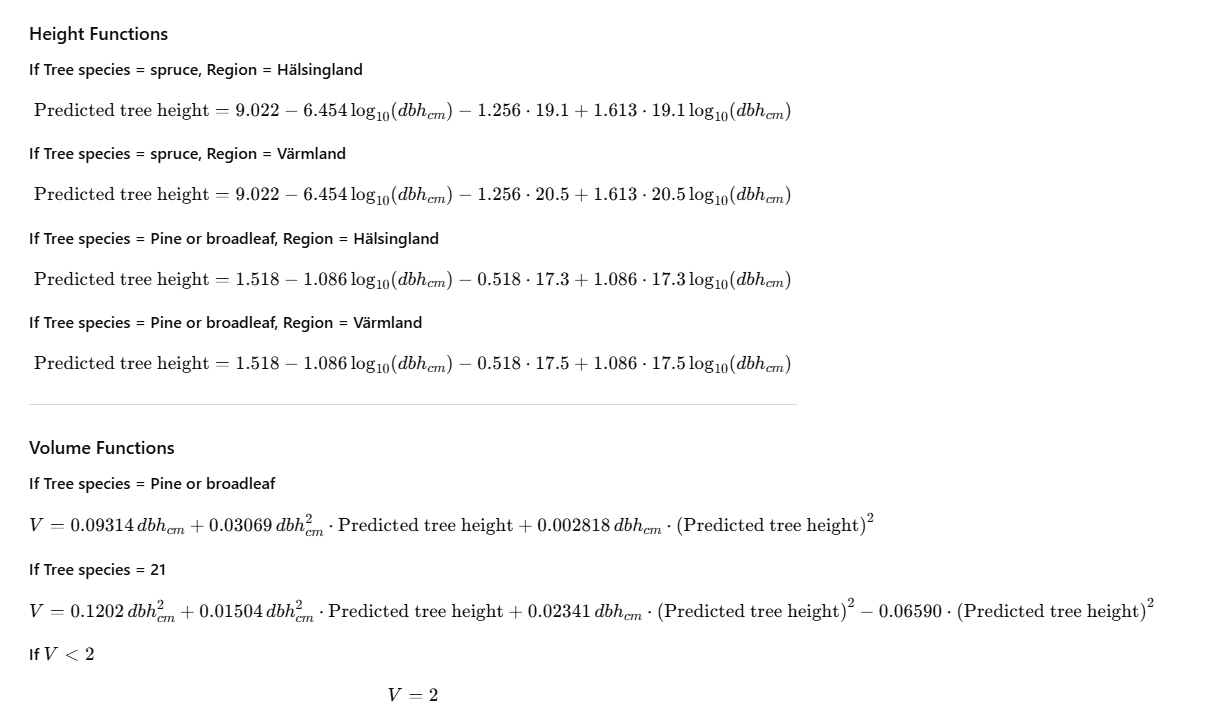

### Species presence and richness modelling

Table S2. The number of sites in each forest management type that had species of each taxa present. 40 was the maximum number of sites surveyed in each forest management type.

| Taxa | Forest Management Type | Number of iItes |
| --- | --- | --- |
| Lichens | Set-aside | 40 |
|  | Retention patch | 40 |
|  | Young forest | 40 |
| Bryophytes | Set-aside | 36 |
|  | Retention patch | 32 |
|  | Young forest | 33 |
| Polypores | Set-aside | 36 |
|  | Retention patch | 29 |
|  | Young forest | 21 |

Table S3. The distribution of environmental variables in each management type across our dataset.

| Management Type | Environment Data | Unit | min | max | mean | sd |
| --- | --- | --- | --- | --- | --- | --- |
| Set-aside | Standing Deadwood | m^3^ per ha | 0 | 39 | 9.5 | 8.1 |
|  | Lying Deadwood | m^3^ per ha | 0 | 190 | 22 | 34 |
|  | Living Trees | m^3^ per ha | 69 | 600 | 280 | 130 |
|  | Living Stem Density | Stems per ha | 390 | 2800 | 1400 | 650 |
| Retention Patch | Standing Deadwood | m^3^ per ha | 0 | 51 | 4.8 | 8.2 |
|  | Lying Deadwood | m^3^ per ha | 0 | 79 | 15 | 17 |
|  | Living Trees | m^3^ per ha | 37 | 390 | 240 | 100 |
|  | Living Stem Density | Stems per ha | 390 | 4800 | 1800 | 1100 |
| Young Forest | Standing Deadwood | m^3^ per ha | 0 | 13 | 0.58 | 2.1 |
|  | Lying Deadwood | m^3^ per ha | 0 | 17 | 3 | 3.8 |
|  | Living Trees | m^3^ per ha | 24 | 220 | 110 | 57 |
|  | Living Stem Density | Stems per ha | 710 | 8300 | 3100 | 1600 |

Table S4. Correlations between each environmental variable. No variables have a correlation of more than 0.5.

|  | Standing Deadwood Volume | Lying Deadwood Volume | Living Tree Volume | Living Tree Density |
| --- | --- | --- | --- | --- |
| Standing Deadwood Volume | - | 0.46 | 0.46 | -0.31 |
| Lying Deadwood Volume | 0.46 | - | 0.31 | -0.14 |
| Living Tree Volume | 0.46 | 0.31 | - | -0.35 |
| Living Tree Density | -0.31 | -0.14 | -0.36 | - |

Table S5. The post hoc test of how, for lichen and bryophyte data, species richness in each forest management type compares. Full model details in **Table 2** (main manuscript). Species Richnes was not significantly related to forest management type in the polypore GLm so we did not undertake post hoc tests.

| Taxa | Contrast | Estimate | Se | Df | z.ratio | p.value |
| --- | --- | --- | --- | --- | --- | --- |
| Lichens | **Young forest – Retention patch** | **-0.450** | **0.0620** | **Inf** | **-7.264** | **<.0001** |
|  | **Young forest – Set-aside** | **-0.564** | **0.1330** | **Inf** | **-7.657** | **<.0001** |
|  | Retention patch – Set-aside | -0.114 | 0.0582 | Inf | -1.952 | 0.12 |
| Bryophytes | Young forest – Retention patch | -0.971 | 1.58 | 105 | -0.615 | 0.8122 |
|  | **Young forest – Set-aside** | **-7.707** | **2.26** | **105** | **-3.410** | **0.0026** |
|  | **Retention patch – Set-aside** | **-6.736** | **1.67** | **105** | **-4.031** | **0.0003** |

Table S6. The distribution of environmental variables between forest management types for sites where bryophytes were found.

| Management Type | Environment Data | Unit | min | max | mean | sd |
| --- | --- | --- | --- | --- | --- | --- |
| Set-aside | Standing Deadwood | m^3^ per ha | 0 | 39 | 10 | 8.1 |
|  | Lying Deadwood | m^3^ per ha | 0.46 | 190 | 24 | 35 |
|  | Living Wood | m^3^ per ha | 110 | 600 | 290 | 130 |
|  | Living Stem Density | Stems per ha | 390 | 2800 | 1300 | 580 |
| Retention Patch | Standing Deadwood | m^3^ per ha | 0 | 51 | 5.7 | 9 |
|  | Lying Deadwood | m^3^ per ha | 1.7 | 79 | 19 | 17 |
|  | Living Wood | m^3^ per ha | 37 | 390 | 250 | 100 |
|  | Living Stem Density | Stems per ha | 390 | 4800 | 1900 | 1200 |
| Young Forest | Standing Deadwood | m^3^ per ha | 0 | 13 | 0.64 | 2.4 |
|  | Lying Deadwood | m^3^ per ha | 0.15 | 17 | 3.6 | 3.9 |
|  | Living Wood | m^3^ per ha | 24 | 210 | 110 | 60 |
|  | Living Stem Density | Stems per ha | 1400 | 8300 | 3200 | 1600 |

Table S7. The relationship between environmental variables and forest management types and the probability of finding at least one species, for bryophytes and polypores. These values were calculated using hurdle models, and the species richness parts of the models are in **Table 2 of the main manuscript**. Standard Error of the model parameters are shown in column SE, and the P values are shown in column Pr.

|  | Environmental Variables | Species Presence Relationship | SE | Pr |
| --- | --- | --- | --- | --- |
| Bryophytes | Standing Deadwood Volume | 1.51 | 1.6 | 0.35 |
|  | **Lying Deadwood Volume** | **8.56** | **2.2** | **<0.001** |
|  | Living Tree Volume | -0.20 | 0.7 | 0.78 |
|  | Living Tree Density | -0.57 | -1.1 | 0.26 |
|  | **Retention Patch** | **-3.14** | **1.4** | **0.03** |
|  | Set aside | -1.46 | 1.45 | 0.31 |
| Polypores | Standing Deadwood Volume | -0.26 | 1.0 | 0.79 |
|  | **Lying Deadwood Volume** | **4.70** | **1.1** | **<0.001** |
|  | Living Tree Volume | -0.06 | 0.5 | 0.90 |
|  | Living Tree Density | -0.59 | 0.3 | 0.54 |
|  | Retention Patch | -0.99 | 0.9 | 0.26 |
|  | Set aside | 1.08 | 1.0 | 0.29 |

Table S8. The distribution of environmental variables between forest management types for sites where polypores were found.

| Management Type | Environment Data | Unit | min | max | mean | sd |
| --- | --- | --- | --- | --- | --- | --- |
| All | Standing Deadwood | m^3^ per ha | 0 | 51 | 6.4 | 8.5 |
|  | Lying Deadwood | m^3^ per ha | 0 | 190 | 18 | 26 |
|  | Living Wood | m^3^ per ha | 27 | 600 | 230 | 130 |
|  | Living Stem Density | Stems per ha | 390 | 8300 | 2000 | 1300 |
| Set-aside | Standing Deadwood | m^3^ per ha | 0 | 39 | 10 | 8.3 |
|  | Lying Deadwood | m^3^ per ha | 0 | 190 | 25 | 35 |
|  | Living Wood | m^3^ per ha | 110 | 600 | 290 | 130 |
|  | Living Stem Density | Stems per ha | 390 | 2800 | 1300 | 590 |
| Retention Patch | Standing Deadwood | m^3^ per ha | 0 | 51 | 5.8 | 9.4 |
|  | Lying Deadwood | m^3^ per ha | 1.2 | 79 | 19 | 18 |
|  | Living Wood | m^3^ per ha | 37 | 390 | 250 | 110 |
|  | Living Stem Density | Stems per ha | 580 | 4400 | 1800 | 970 |
| Young Forest | Standing Deadwood | m^3^ per ha | 0 | 13 | 0.99 | 2.9 |
|  | Lying Deadwood | m^3^ per ha | 0.62 | 17 | 4.8 | 4.5 |
|  | Living Wood | m^3^ per ha | 27 | 210 | 110 | 54 |
|  | Living Stem Density | Stems per ha | 1500 | 8300 | 3300 | 1700 |

### Dissimilarity modelling

#### Lichen Generalised Dissimilarity Modelling Plots

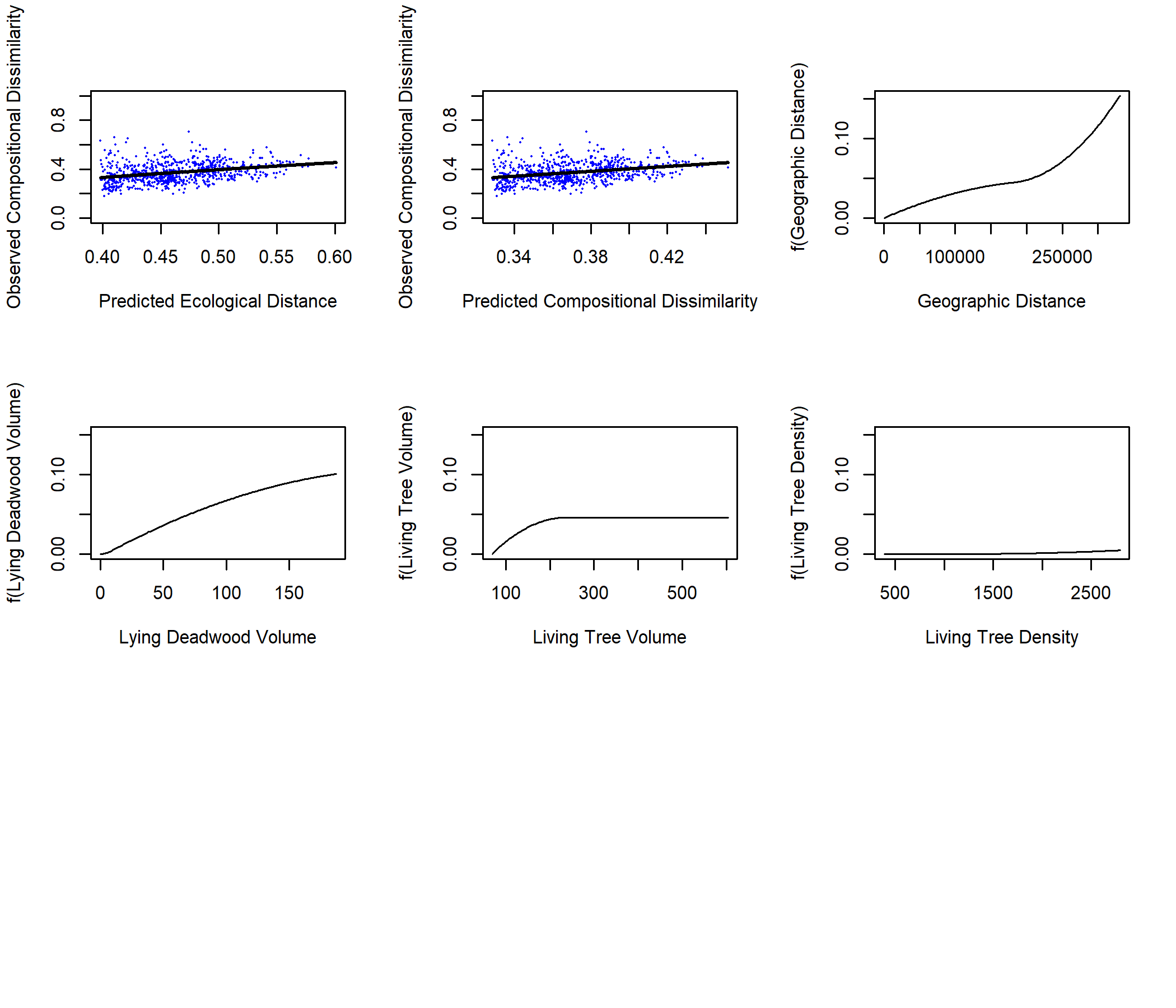

Figure S1. Plots from the Generalised Dissimilarity Modelling (GDM) of lichen beta diversity in Set-asides against geographic and environmental distance. (a) Observed dissimilarity as a function of GDM-predicted ecological distance, where each point represents a site-pair, and the line represents the GDM predicted dissimilarity. (b) Observed dissimilarity as a function of GDM predicted dissimilarity, with the line of equality provided. This GDM includes data from all 780 lichen Set-aside site-pairs, using four environmental predictors (Table 1), explaining 9.5% of the variation in observed dissimilarities and with an intercept of 0.40.

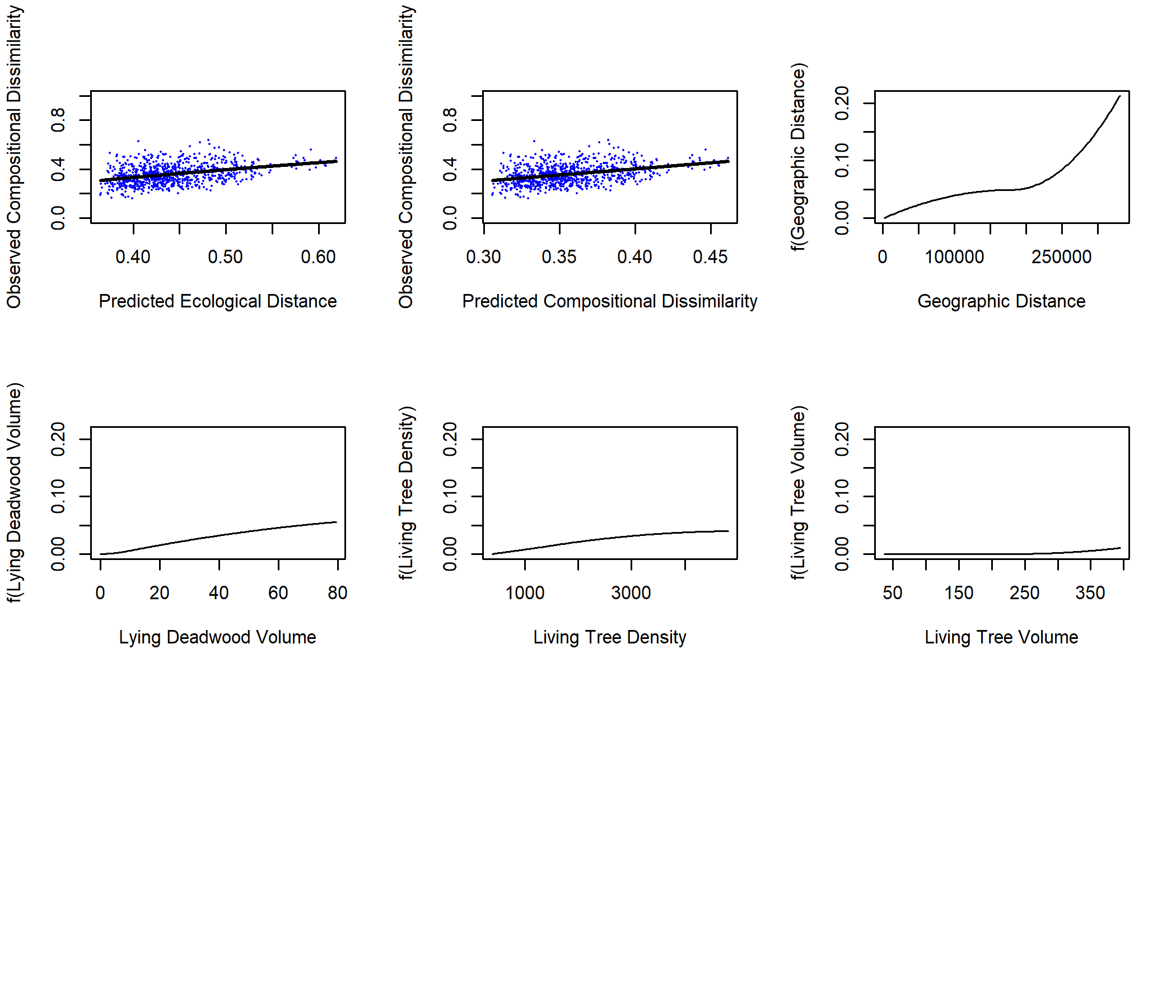

Figure S2. Plots from the Generalised Dissimilarity Modelling (GDM) of lichen beta diversity in Retention Patches against geographic and environmental distance. (a) Observed dissimilarity as a function of GDM-predicted ecological distance, where each point represents a site-pair, and the line represents the GDM predicted dissimilarity. (b) Observed dissimilarity as a function of GDM predicted dissimilarity, with the line of equality provided. This GDM includes data from all 780 lichen Retention Patch site-pairs, using four environmental predictors (Table 1), explaining 11.1% of the variation in observed dissimilarities and with an intercept of 0.34.

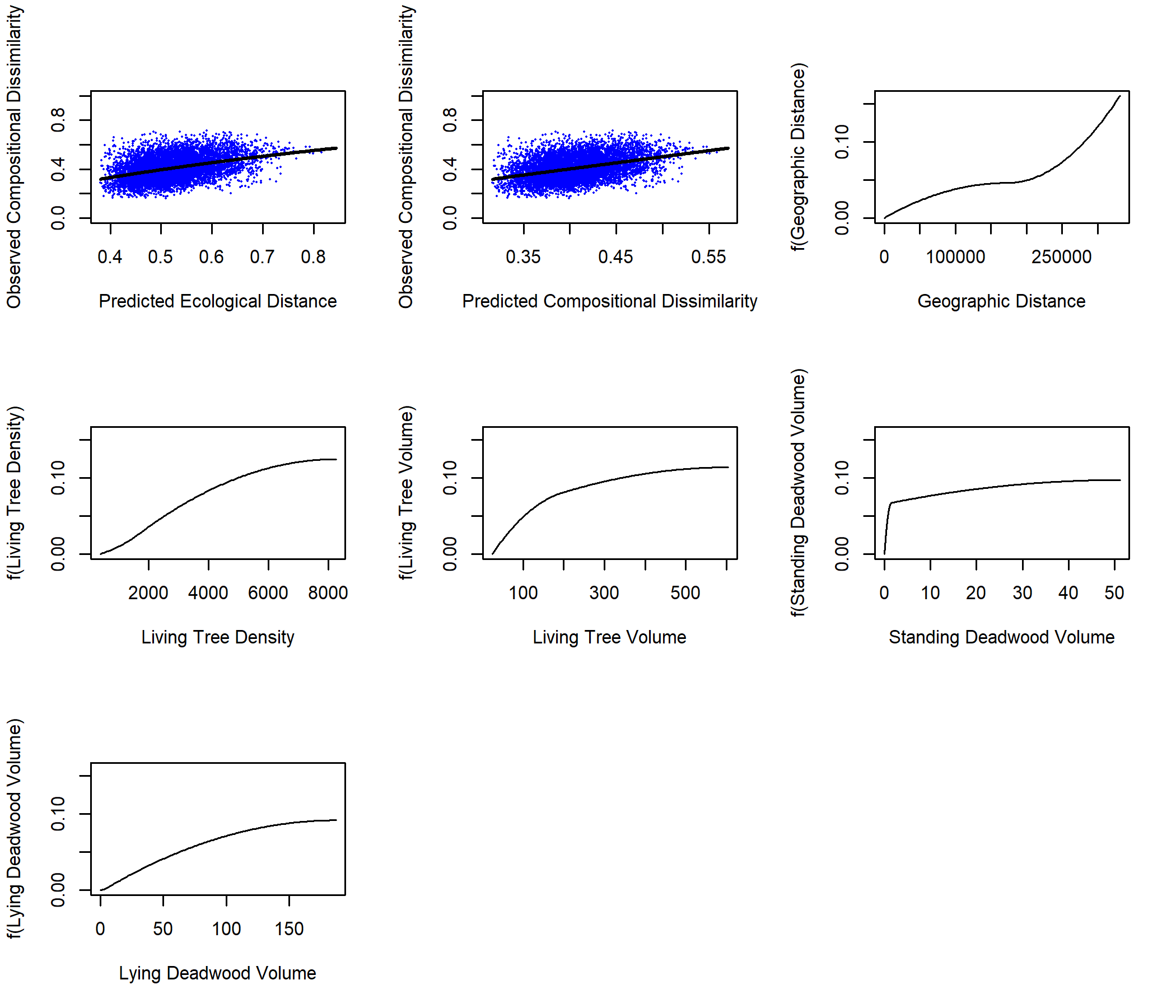

Figure S3. Plots from the Generalised Dissimilarity Modelling (GDM) of lichen beta diversity in Young Forests against geographic and environmental distance. (a) Observed dissimilarity as a function of GDM-predicted ecological distance, where each point represents a site-pair, and the line represents the GDM predicted dissimilarity. (b) Observed dissimilarity as a function of GDM predicted dissimilarity, with the line of equality provided. This GDM includes data from all 780 lichen Young Forests site-pairs, using four environmental predictors (Table 1), explaining 30.0% of the variation in observed dissimilarities and with an intercept of 0.30.

#### Bryophyte Generalised Dissimilarity Modelling Plots

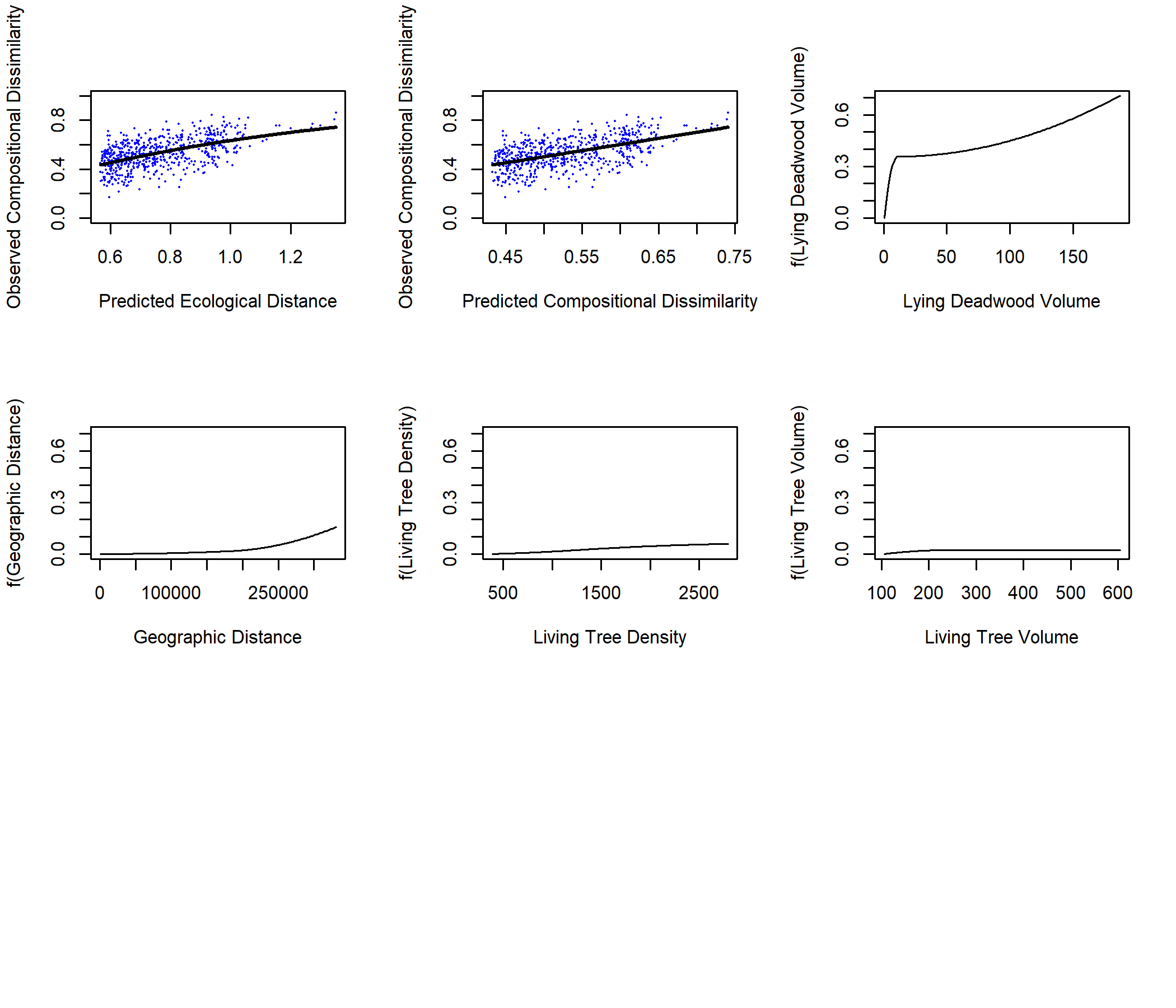

Figure S4. Plots from the Generalised Dissimilarity Modelling (GDM) of bryophyte beta diversity in Set-asides against geographic and environmental distance. (a) Observed dissimilarity as a function of GDM-predicted ecological distance, where each point represents a site-pair, and the line represents the GDM predicted dissimilarity. (b) Observed dissimilarity as a function of GDM predicted dissimilarity, with the line of equality provided. This GDM includes data from the 36 Set-aside sites with bryophytes present, meaning 630 site-pairs, using four environmental predictors (Table 1), and explaining 28.8% of the variation in observed dissimilarities and with an intercept of 0.56.

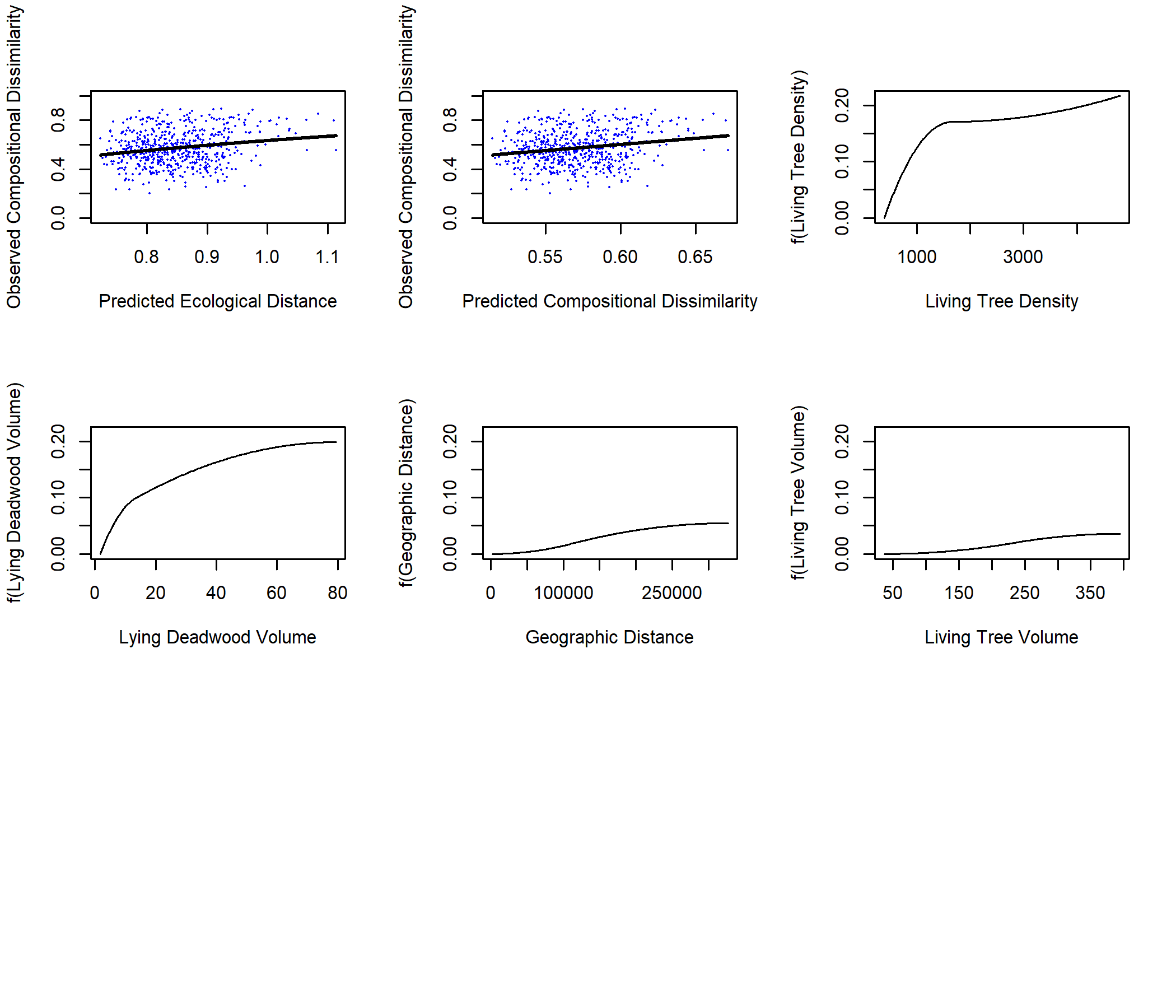

Figure S5. Plots from the Generalised Dissimilarity Modelling (GDM) of bryophyte beta diversity in Retention Patches against geographic and environmental distance. (a) Observed dissimilarity as a function of GDM-predicted ecological distance, where each point represents a site-pair, and the line represents the GDM predicted dissimilarity. (b) Observed dissimilarity as a function of GDM predicted dissimilarity, with the line of equality provided. This GDM includes data from the 32 Retention Patch sites with bryophytes present, meaning 496 site-pairs, using four environmental predictors (Table 1), and explaining 3.6% of the variation in observed dissimilarities and with an intercept of 0.69.

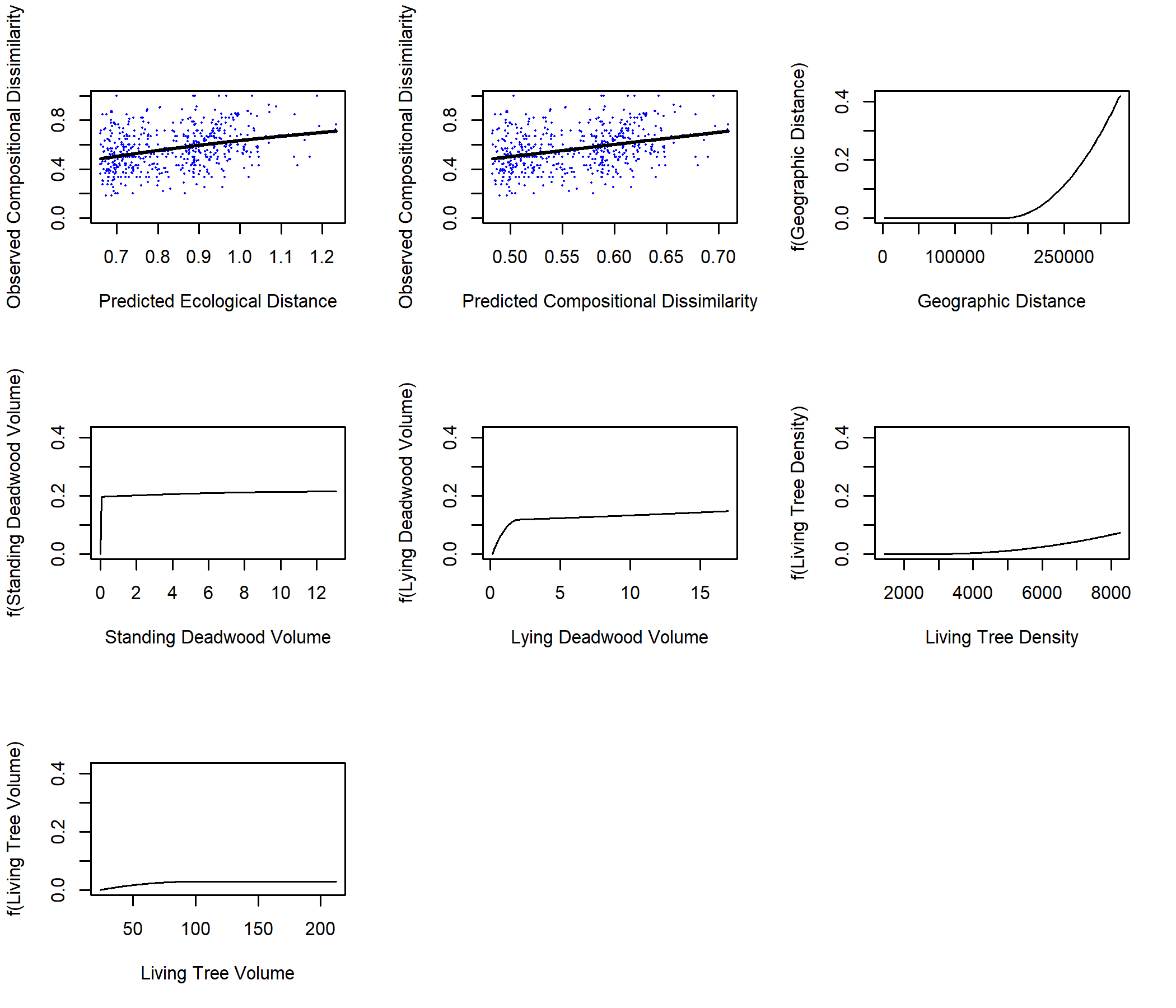

Figure S6. Plots from the Generalised Dissimilarity Modelling (GDM) of bryophyte beta diversity in Young Forests against geographic and environmental distance. (a) Observed dissimilarity as a function of GDM-predicted ecological distance, where each point represents a site-pair, and the line represents the GDM predicted dissimilarity. (b) Observed dissimilarity as a function of GDM predicted dissimilarity, with the line of equality provided. This GDM includes data from the 33 Young Forest sites with bryophytes present, meaning 528 site-pairs, using four environmental predictors (Table 1), and explaining 9.2% of the variation in observed dissimilarities and with an intercept of 0.66.

#### Polypore Generalised Dissimilarity Modelling Plots

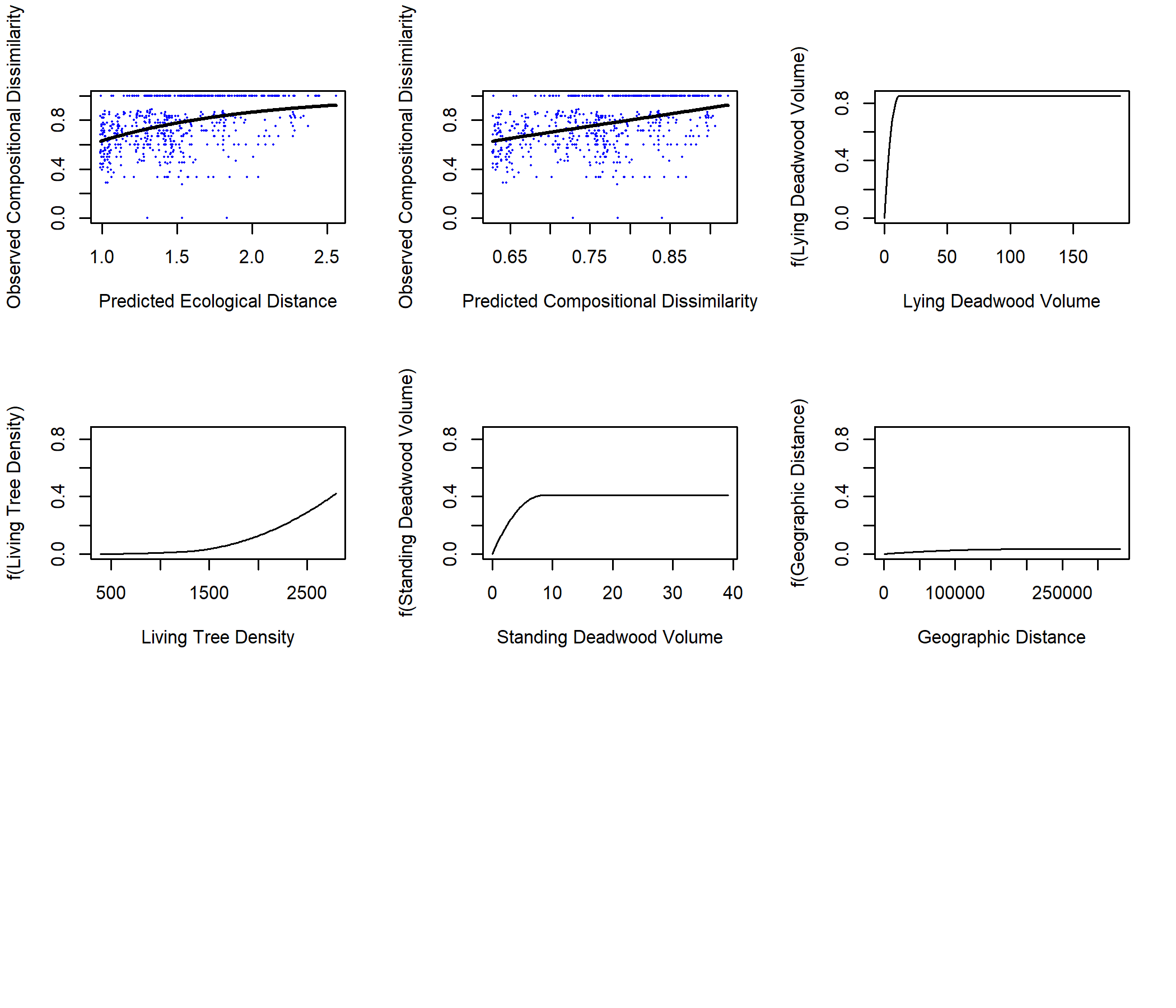

Figure S7. Plots from the Generalised Dissimilarity Modelling (GDM) of polypore beta diversity in Set-asides against geographic and environmental distance. (a) Observed dissimilarity as a function of GDM-predicted ecological distance, where each point represents a site-pair, and the line represents the GDM predicted dissimilarity. (b) Observed dissimilarity as a function of GDM predicted dissimilarity, with the line of equality provided. This GDM includes data from the 36 Set-asides with polypores present, leading to 630 site-pairs, using four environmental predictors (Table 1), and explaining 12.5% of the variation in observed dissimilarities and with an intercept of 0.98.

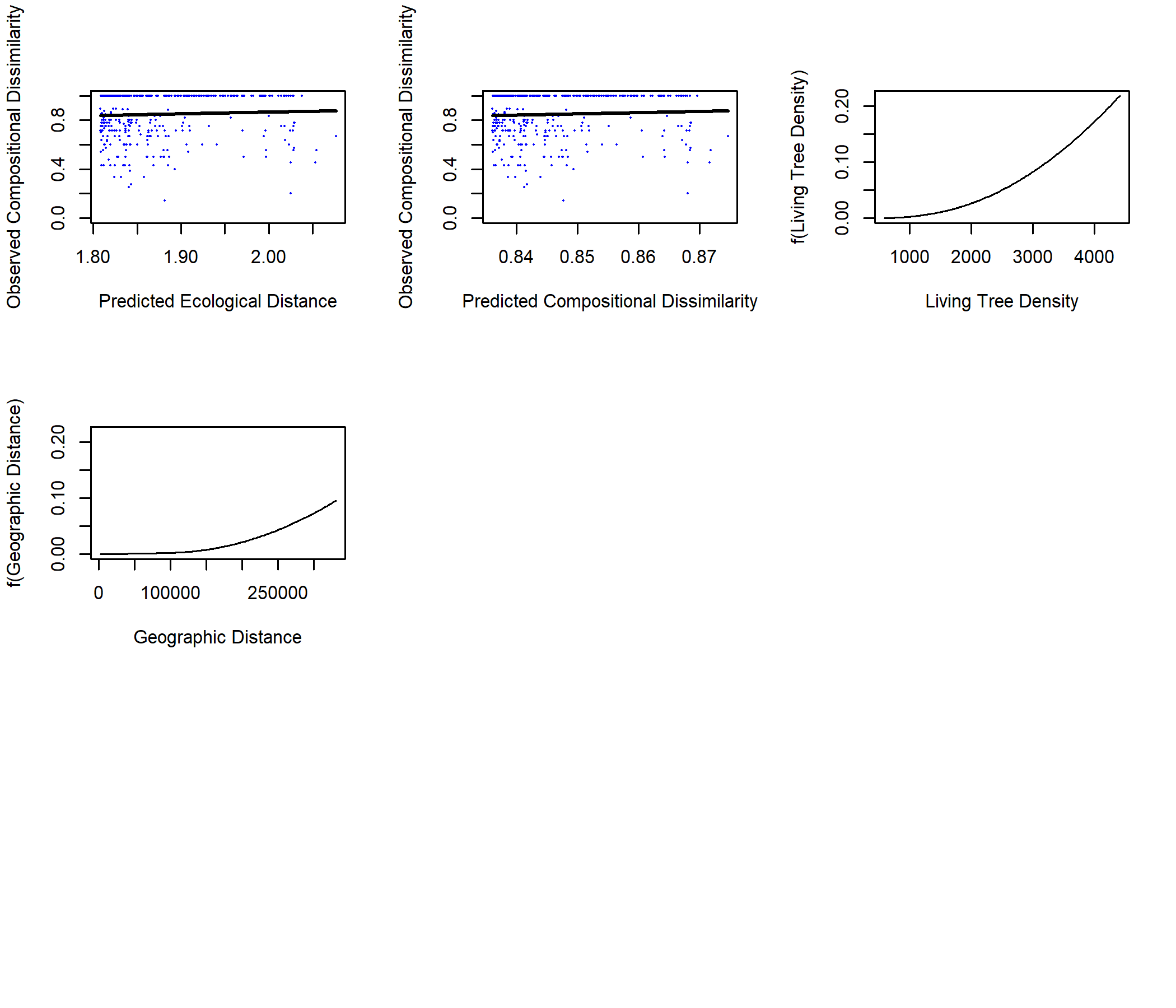

Figure S8. Plots from the Generalised Dissimilarity Modelling (GDM) of polypores beta diversity in Retention Patches against geographic and environmental distance. (a) Observed dissimilarity as a function of GDM-predicted ecological distance, where each point represents a site-pair, and the line represents the GDM predicted dissimilarity. (b) Observed dissimilarity as a function of GDM predicted dissimilarity, with the line of equality provided. This GDM includes data from the 29 Retention Patches were polypores were found, so lichen 406 site-pairs, using four environmental predictors (Table 1), explaining 0.23% of the variation in observed dissimilarities and with an intercept of 1.8.

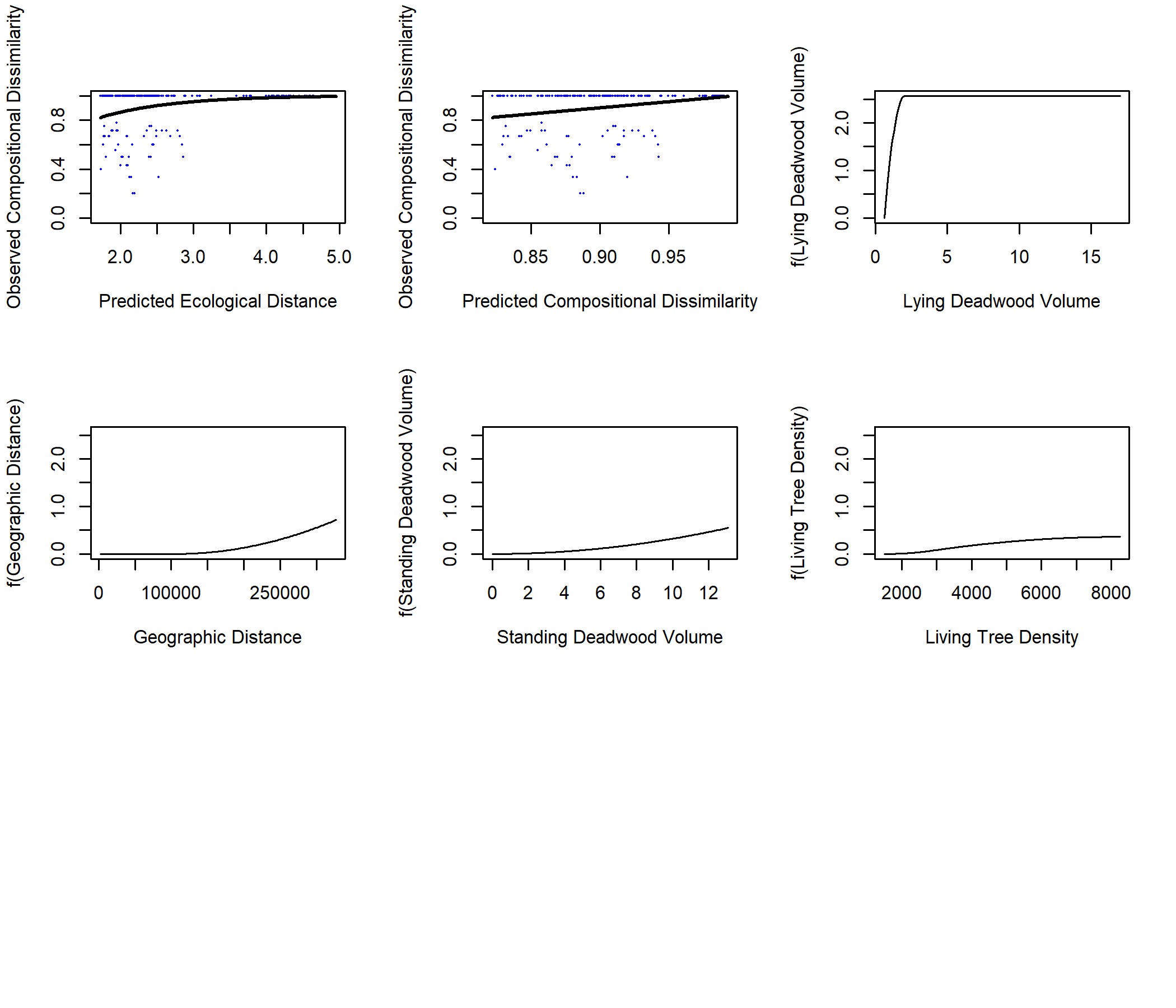

Figure S9. Plots from the Generalised Dissimilarity Modelling (GDM) of polypores beta diversity in Young Forests against geographic and environmental distance. (a) Observed dissimilarity as a function of GDM-predicted ecological distance, where each point represents a site-pair, and the line represents the GDM predicted dissimilarity. (b) Observed dissimilarity as a function of GDM predicted dissimilarity, with the line of equality provided. This GDM includes data from the 21 Young Forests were polypores were found, so lichen 210 site-pairs, using four environmental predictors (Table 1), explaining 10.0% of the variation in observed dissimilarities and had an intercept of 1.7.
